# Human meibomian gland organoids to study epithelial homeostasis and dysfunction

**DOI:** 10.1101/2025.06.10.658800

**Authors:** Chuyue Yu, Xichen Wan, Jinsong Wei, Xingru Wu, Zhaoting Xu, Xiaoye Wang, Yabo Mi, Yiming Zhang, Dan Wu, Xujiao Zhou, Qihua Le, Jianjiang Xu, Chen Zhao, Xinghuai Sun, Xingtao Zhou, Jiaxu Hong, Bing Zhao

**Author notes:** Correspondence and requests for materials should be addressed to J.H. or B.Z. (;. These authors contributed equally to this manuscript (Chuyue Yu, Xichen Wan, Jinsong Wei).

## Abstract

Meibomian glands (MGs) are holocrine glands that secrete lipids to maintain the homeostasis of ocular surface, and their dysfunction leads to dry eye disease. Herein, we established long-term 3D organoid culture for murine and human MGs, which retained the cell lineages and lipid-producing ability. The organoids mimicked the drug treatment responses and generated functional MGs after orthotopic transplantation. Inspired by organoid cultures, we found FGF10 eye drops could rescue all-trans retinoic acid-induced MG dysfunction in mice. Besides, nicotinamide uniquely hampered the human MG organoid expansion by inhibiting FGF10 signaling. Single-cell atlas and lipidome not only aligned the delineated cell types and featured lipids between human MGs and organoids, but highlighting MAPK signaling inhibition enhanced acinar cell differentiation and functional maturation of MG organoids. In summary, this study established an organoid platform to explore epithelial homeostasis and dysfunction of MGs, facilitating drug development and regenerative medicine for dry eye disease.

## Main

Meibomian glands (MGs), the holocrine sebaceous glands located in the eyelids, are essential for the homeostasis of the tear film and ocular surface. MGs are featured by a central duct surrounded by acini that responsible for lipid production. Under homeostasis, the basal cells in acini could self-renew and differentiate to acinar cells, which is also called meibocytes. The meibocytes begin to synthesize lipid while migrating towards the center of acini. Finally, the meibocytes rupture to release the lipid content. Delivered by the central duct to the eyelid margin, the secreted lipid forms outermost layer of the preocular tear film to prevent tear evaporation and environmental insults ^1^.

Meibomian gland dysfunction (MGD) is the leading cause of evaporative dry eye disease ^2^, which is featured by poor vision quality, eye irritation, ocular surface inflammation and discomfort, affecting 38–68% population worldwide. However, the only treatment for MGD is palliative care ^3^. Various risky factors were reported, including aging, drug treatment and congenital defect ^4–11^, while underlying pathology of MGD would be the stem cell exhaustion. To this end, the culture of MG stem cells, which have could self-renew and differentiate into functional meibocytes *in vitro*, would be promising for the MGD therapeutics. However, existing 2D immortalized MG epithelial cells are difficult to simulate in *in vivo* responses ^12, 13^. Thus, it is urgent to create an *ex vivo* model for exploring the homeostasis and dysfunction of MGs.

Organoids with three-dimensional (3D) structures derived from adult stem cells accurately mimic physiological aspects of parental tissues *ex vivo*. Thus, organoids are used to model organ development and disease progression ^14^. However, no approach has been established to generate human MG organoids yet, limiting the study of the physiology and pathology of MGs. In this study, we achieved long-term expansion of murine and human MGs as 3D organoids, which recapitulated the characteristics of tissues, and modeled the MG degenerative diseases induced by chemical injuries *ex vivo*. Notably, human MG organoids could engraft and undergo functional maturation after orthotopic transplantation into mice. Further, we probed the cell types and lipidome of parallel human MG tissues and organoids using scRNA-seq and lipidomic analysis, highlighting the MAPK signaling is critical for meibocyte maturation. Taken together, this study described an organoid platform to explore the homeostasis and diseases of MGs and provided a useful platform for drug testing and regenerative medicine.

## Results

### Establishment of functional murine meibomian gland organoids

Before formulating the culture protocol for meibomian gland (MG) organoids, we firstly examined the key signaling pathways in the meibomian gland tissues (Fig. 1A). FGF signaling is essential for mammary gland development, and loss of Fgf10 or its receptor Fgfr2 led to placodes defect during embryogenesis ^15, 16^. The Wnt signaling is the key driver of adult stem cells in most tissues. As the receptor for R-spondins, Lgr5 marks adult stem cells in actively self-renewing organs ^17^. In addition to the extracellular signals, nicotinamide phosphoribosyltransferase (Nampt) is key enzyme that regulates intracellular nicotinamide adenine dinucleotide (NAD) pool, which is crucial for cellular redox stage ^18^. Immunostaining showed that Fgfr2, Lgr5 and Nampt were highly expressed in meibomian gland tissues, which were marked by Krt14 (Fig. 1B). Therefore, we supplemented FGF10, R-spondin-1 and nicotinamide as the core factors in murine MG organoid culture medium. During 9 days’ incubation, the seeded MG cell gradually formed the spheroid structure, and became solid and opaque during expansion. (Fig. 1C and Extended Data Fig. 1A). H&E staining showed organoids and tissues shared similar architecture: the outer layer of organoid was lined with small and compact cells, which resembled the basal cells. While the internal cells displayed lower nucleocytoplasmic ratios and condensed cell nuclei ruptures, which resembled the differentiated acinar cells (Fig. 1D).

**Figure 1.**
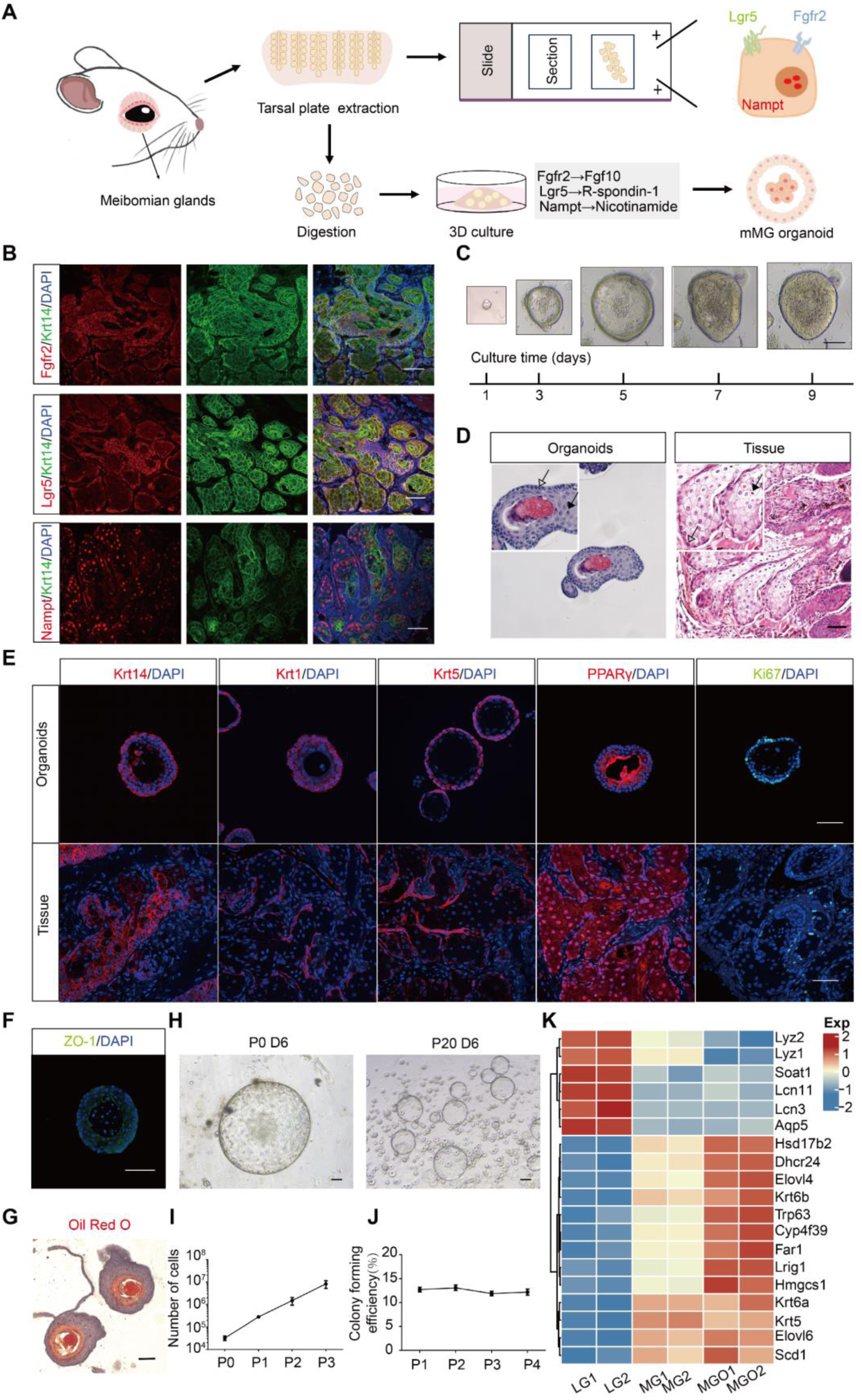
Generation of functional murine MG organoids. (A) Schematic diagram of organoid culture from murine MG tissue. (B) Immunofluorescent staining for Fgfr2, Lgr5 and Nampt in murine MG tissue (marked by Krt14). Scale bar, 50 μm. (C) Representative bright-field images of the outgrowth of single organoid during the 9-days culture period. Scale bar, 100 μm. (D) H&E staining of passage 5, day 9 organoid and the murine MG tissue. White box indicates regions shown in higher magnifications (x2). Filled arrow indicate basal cells, and empty arrow indicate meibocytes with pyknotic nucleus. Scale bar, 50 μm. (E) Immunofluorescent staining for the MG epithelial cell marker (Krt14), ductal cell marker (Krt1), basal cell marker (Krt5), meibocyte marker (PPARγ) and the proliferation marker Ki67 in organoids and in tissue. Scale bar, 50 μm. (F) Immunofluorescent staining for ZO-1 in murine MG organoids. Scale bar, 50 μm. (G) Oil Red O staining of lipids in murine MG organoids. Scale bar, 50 μm. (H) Representative images of MG organoids outgrowth from primary organoids to passage 20. Scale bar, 100 μm. P: passage; D: day (I-J) The cell number (I) and colony forming efficiency (J) of MG organoids during passage 0-4. (K) Heatmap indicating the expression of MG markers and lipid synthesis related genes in murine MG tissues and organoids. 2 independent mice for each group. LG, lacrimal gland; MG, meibomian gland; MGO, meibomian gland organoid. See also Figure S1

To further confirm the cell lineages in organoids, we stained the typical cell markers. In the MGs, the basal acinar cells (Krt5^+^) retained the ability of self-renewal, and moved toward the center of the acini to form terminal differentiated gland acinar cells/ meibocytes (PPARγ^+^). Ideally, the organoids well preserved the cell lineages and their niche in MGs, exhibiting proliferative Ki67^+^ or Krt5^+^ basal acinar cells lining the out layers, and PPARγ^+^ meibocytes locating in the inner layers (Fig. 1E). Besides, the inner cell also gained cell polarity, with the apical side facing inside (Fig. 1F). Importantly, the MG organoids could be long-term expanded up to 20 passages (Fig. 1H, I and J).

The main function of MG is meibum/lipid secretion. Oil Red O staining of MG organoids showed that inner layer cells were Oil Red O positive, and the inner cavity displayed much condense staining (Fig. 1G, Extended Data Fig. 1B), indicating the meibocytes secreted lipid into the cavity. To further characterized the transcriptomic similarity between MGs and MG organoids, we performed bulk RNA-seq, and set irrelevant lacrimal gland as controls. The organoids displayed the similar gene expression pattern with the MGs (Extended Data Fig. 1C, D), and showed a significant enrichment of MG signatures: *Krt6*, *Lrig1* and lipid synthesis-related genes *ElovI6*, *Scd1*, *Far1* and *Hmgcs1* (Fig. 1K and Extended Data Fig. 1D). Besides, we found that age and sex do not affect the MGO establishment (Extended Data Fig. 1E). In summary, the long-term expandable and functional MG organoids preserved the cell lineages, gene expression signatures and lipid-secretion capability of MGs.

### FGF10 sustained MG organoids expansion and prevented MGs dysfunction

The regeneration of MG depends on the self-renewable acinar basal cells, and the long-term expandable MG organoid culture system highlighted critical signaling pathways that boost basal cell amplification (Fig. 1E). We reasoned that the dominating signaling could be identified using organoid culture system. After removing FGF10, Noggin and R-spondin1 from culture medium individually, we found that MG organoids were most sensitive to FGF10, rather than Noggin or R-spondin1 (Fig. 2A-C). FGF10 was critical factor in regulating the development of eyelid and the proliferation of immortalized human MG epithelial cells ^19, 20^, while its role in regeneration was less investigated Inspired by above, we then determined the function of FGF10 in MG dysfunction (MGD).

**Figure 2.**
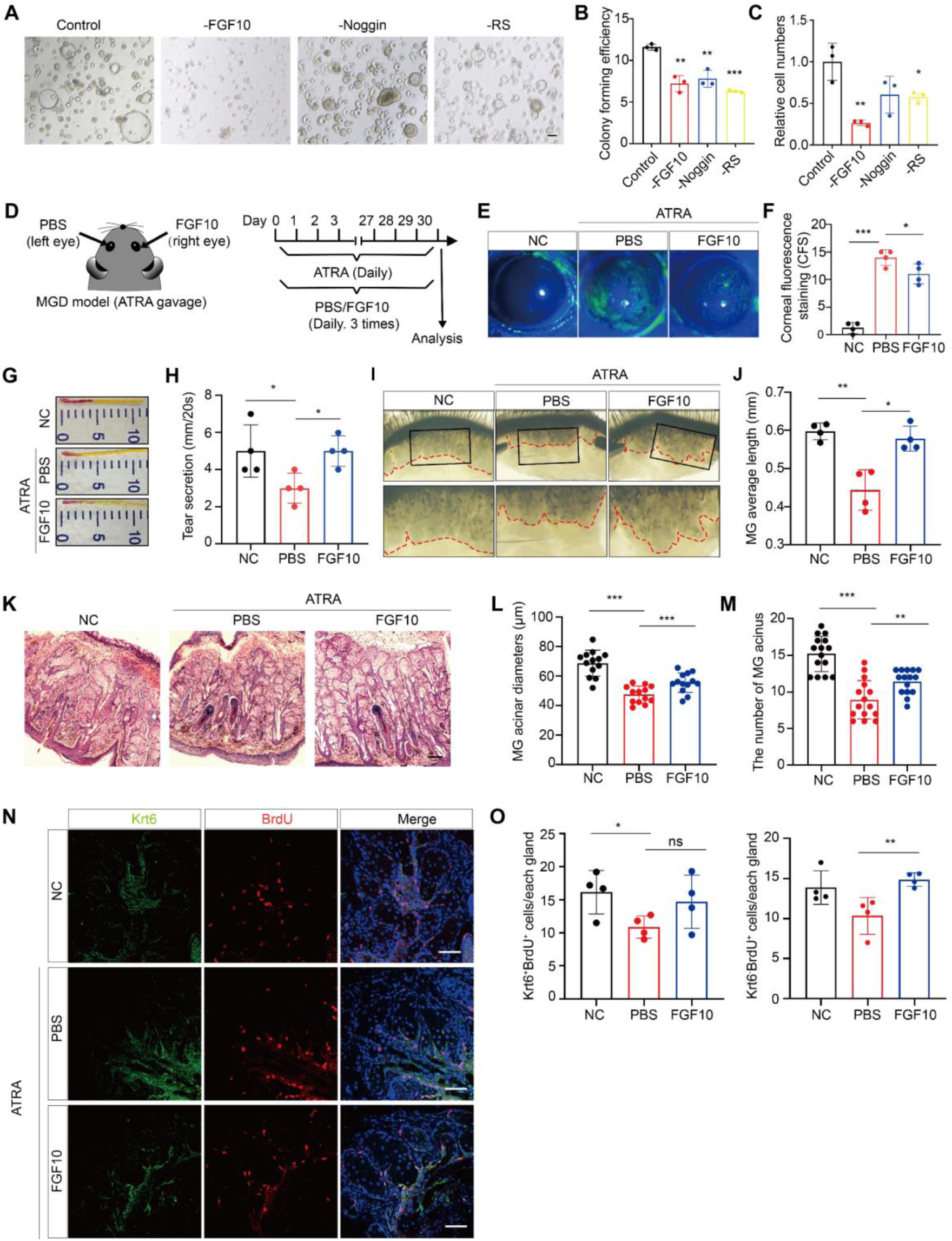
FGF10 is required for the MG organoids expansion and prevents MG disfunction. (A) Representative bright-field images of the MG organoids in 4 conditions (removing the FGF10, Noggin, Rspo-1 and control). Scale bar, 100 μm. (B-C) Quantification of colony forming efficiency (B) and cell numbers (C) of organoids in 4 indicated conditions. Data were represented as mean ± SEM in 3 independent experiments. Unpaired two-tailed Student’s *t* test: \**p*<0.05, ** *p*<0.01, \*\*\**p*<0.001. (D) Schematic of the workflow and the timeline for ATRA-induced MGD model and FGF10 eye drops administration. (E-F) Corneal fluorescein staining (E) images and (F) scores of normal control mice and ATRA-induced MGD mice treated with PBS or FGF10 eye drops. NC: normal control group without any treatment; PBS: ATRA-induced MGD group treated with PBS eye drops; FGF10: ATRA-induced MGD group treated with FGF10 eye drops. Data were represented as mean ± SEM. n=4 mice for each group. Unpaired two-tailed Student’s *t* test: \**p*<0.05, \*\*\**p*<0.001. (G-H) Representative images (G) and measurement (H) of tear secretion volume using phenol red thread test of indicated groups. Data were represented as mean ± SEM. n=4 mice for each group. Unpaired two-tailed Student’s *t* test: \**p*<0.05. (I-J) Representative bright-field images of MGs (I) and quantification of MGs length (J) of indicated groups. Red dotted line indicates margin of MGs. Lower panel: higher magnifications of upper panel. Data were represented as mean ± SEM. n=4 mice for each group. Unpaired two-tailed Student’s *t* test: \**p*<0.05, ** *p*<0.01. (K-M) H&E staining (K) and quantification of size (L) and number (M) of MG acini of indicated groups. Quantified data were determined in 12 randomly selected fields. Data were represented as mean ± SEM. Unpaired two-tailed Student’s *t* test: ** *p*<0.01, *** *p*<0.001. Scale bar, 100μm. (N) Immunofluorescent staining for Krt6 (ductal cell marker) and BrdU (proliferation marker) of indicated groups. Scale bar, 50μm. (O) Quantification of number of proliferative non-ductal cells (Krt6^-^/BrdU^+^ cells) and ductal cells (Krt6^+^/BrdU^+^ cells) in indicated groups. Data were represented as mean ± SEM. n=4 mice for each group. Unpaired two-tailed Student’s *t* test: \**p*<0.05, ** *p*<0.01. See also Figure S2 and S3

To established mouse MGD model, we optimized previously described retinoic acid method by replacing 13-*cis* retinoic acid (13-cis-RA) with all-*trans* retinoic acid (ATRA) ^21^, since it is generally accepted that the effects of 13-cis-RA are mediated by isomerization to the more transcriptionally active ATRA form ^22^. Mice were treated with ATRA through gavage for a month to induce MGs damage, and the FGF10 solution was dripped into right eye (PBS for left eye) three times/day as therapeutic group (Fig. 2D). Ophthalmologic examinations, including ocular epithelial integrity, tear secretion and MG length, were conducted to score the MGD development and FGF10 treatment. Compared with untreated group, ATRA+PBS group showed pervaded corneal fluorescein staining (CFS), indicating compromised ocular epithelium. In addition, tear secretion ability and MG length were also reduced in ATRA+PBS group (Fig. 2E-J). Histologically, ATRA+PBS also reduced the number and size of the acini in the MGs (Fig. 2K-M) and inhibited the BrdU^+^ cells in both the ductal (Krt6^+^ cells) and non-ductal cells (Krt6^-^ cells) of the MGs (Krt6^+^ cells) (Fig. 2N). Although no significant effects were found in the expression of PPARγ, the lipid-synthesis (oil-red) was inhibited in ATRA+PBS group (Extended Data Fig. 2C-D). Above data suggested ATRA induced MGD model was successfully established. Interestingly, based on the MGD model, ATRA+FGF10 group exhibited significant MG recovery in all aspects, compared with ATRA+PBS group. Ideally, ATRA+FGF10 rescued ocular epithelium integrity, tear secretion, gland architecture, cell proliferation and lipid synthesis.

To dive deeper into the role of FGF10 signaling during MGD, we performed phospho-p38 (downstream indicator of FGF10 signaling) and KRT14 co-immunostaining, and found ATRA treatment reduced the phospho-p38 signal and FGF10 application restores it (Extended Data Fig. 3). These data demonstrated that FGF10 contributed to the MGs homeostasis, and the activation of FGF10 signaling alleviated the progression of MGD.

### MG organoids mimicked the drug responses

Since the MG organoid could recapitulate the physiological aspects of MGs, we reasoned that the MG organoid might be used to discover the drugs that resolved dry eye disease. To demonstrate this, we selected drugs with known effects on MGs: ATRA and 13-cis-RA were confirmed to have a detrimental effect, resulting in cell death, atrophy, and altered gene expression ^23^. Azithromycin (AZM) can stimulate lipid synthesis in iHMGECs and is the prescription medication for the treatment of MGD ^24, 25^. Rapamycin, the mTOR signaling pathway inhibitor, is a potential drug for the treatment of age-related disease ^26^. Dihydrotestosterone (DHT) is a major androgen that has been considered to regulate MG function by suppressing inflammation ^27^, and enhancing the lipids synthesis ^21^.

We exposed the MG organoids to those drugs individually and followed by the morphologic and functional characterizations. Compared to the dimethylsulfoxide (DMSO) group, treatment with 100 μM 13-cis RA, 10 μM Rapamycin, and 10 μM DHT resulted in a significant decrease in organoids size (Fig. 3A, B). The inner cavity of organoids became condense after being treated with 10 μM AZM and 10 μM DHT, suggesting that AZM and DHT might induce the differentiation (Figure 3A). Administration of 100 μM ATRA and 100 μM 13-cis-RA significantly inhibited the expression of proliferation and stemness genes ^12^, including *Krt5*, *Lrig1*, and *Pcna*, as well as protein level of PCNA and Sox2 (Fig. 3C, G). Furthermore, the MG markers, *Krt10* and *Krt14,* decreased after the treatment with ATRA and 13-cis-RA, while the ductal cells marker gene *Krt6* were upregulated. The data suggested that the effects of the ATRA and 13-cis RA on the acinar and ducts cells of the MGs might be different (Extended Data Fig. 3G and Extended Data Fig. 4A, B).

**Figure 3.**
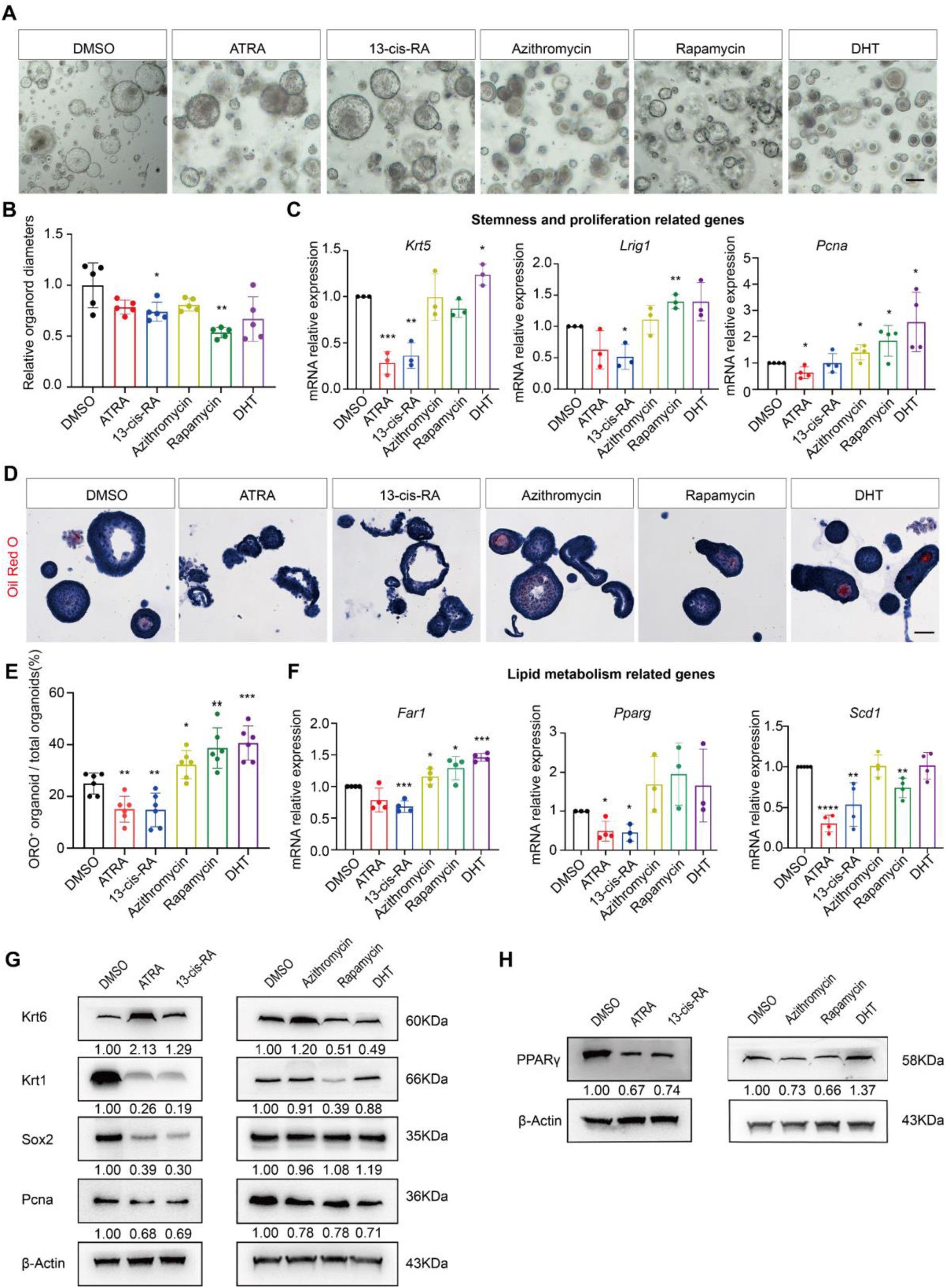
Organoids mimic the drug response of MGs. (A) Representative bright-field images of MG organoids incubated with 0.1%DMSO, 100μM ATRA, 100μM 13-cis RA, 10μM AZM, 10μM Rapamycin and 10μM DHT for 3 days, respectively. Scale bar, 100μm. (B) Quantification of diameters of organoids of indicated treatment. Quantified data were determined in 5 randomly selected fields. Data were represented as mean ± SEM. Unpaired two-tailed Student’s *t* test: \**p*<0.05, ** *p*<0.01. (C) Quantitative real time PCR (qRT-PCR) analysis of the expression level of proliferative and stemness-related genes including *Krt-5, Lrig1* and *Pcna* of the organoids after indicated treatment. Data are represented as mean ± SEM in ≥3 independent experiments. Unpaired two-tailed Student’s *t* test: \**p*<0.05, ** *p*<0.01, \*\*\**p*<0.001. (D) Oil Red O staining of lipids in organoids with indicated treatment. Scale bar, 50 μm. (E) Quantification of the number of Oil Red O ^+^ organoids divided by the total number of organoids with indicated treatment. Quantified data were determined in 6 randomly selected fields. Data are represented as mean ± SEM. Unpaired two-tailed Student’s *t* test: \**p*<0.05, ** *p*<0.01, \*\*\**p*<0.001. (F) Quantitative real time (RT)-PCR analysis of the expression level of lipid metabolism related genes including *Far*, *Pparg* and *Scd1* with RNA extracted from the organoids after indicated treatment. Data were represented as mean ± SEM in ≥3 independent experiments. Unpaired two-tailed Student’s *t* test: \**p*<0.05, ** *p*<0.01, \*\*\**p*<0.001 (G-H) Western blotting analysis of Krt6, Krt1, Sox2, Pcna (stemness-related protein) (G) and PPARγ (metabolism related protein) (H) protein expression in organoids with indicated treatment. β-actin served as a loading control. The relative band intensity that normalized to loading control was marked under each band. n=3 biological replicates See also Figure S4

Regarding to lipid synthesis, AZM, Rapamycin, and DHT treatments resulted in a significant increase in lipid production capacity, while ATRA and 13-cis RA led to a decrease, as indicated by the Oil Red O staining (Fig. 3D, E). Additionally, ATRA and 13-cis RA led to reduction of lipid synthesis-related genes *Far1*, *Pparg* and *Scd1* while promoted fatty acid synthase (*Fasn*) and Perilipin (*Plin2*) expression, suggesting that ATRA and 13-cis RA affected specific stages of differentiation (Fig. 3F and Extended Data Fig. 3C, D). In summary, the MG organoids could rapidly assess the drug effects, and precisely simulated drug responses *ex vivo*.

### Establishment of human MG organoids for orthotopic transplantation

Next, we attempted to establish human MG organoids (hMGO). Refer to murine organoid, we also mapped the expression pattern of FGFR2, LGR5 and NAMPT in human MG tissue (Fig. 4A). Unexpectedly, NAMPT was barely expressed in human MG tissue (Fig. 4B), so we suspected that its substance-nicotinamide might not be required for the human organoid culture. Indeed, the mMGO expansion medium could not support hMGO expansion for more than 5 days. However, once nicotinamide was removed, the hMGO dramatically expanded, so we defined this formula as hMGO expansion medium (Fig. 4C). As expected, the hMGO medium could support long-term expansion of hMGO to 20 passages (P20), as well as nearly 10 times cell amplification per passage (Fig.4D-E).

**Figure 4.**
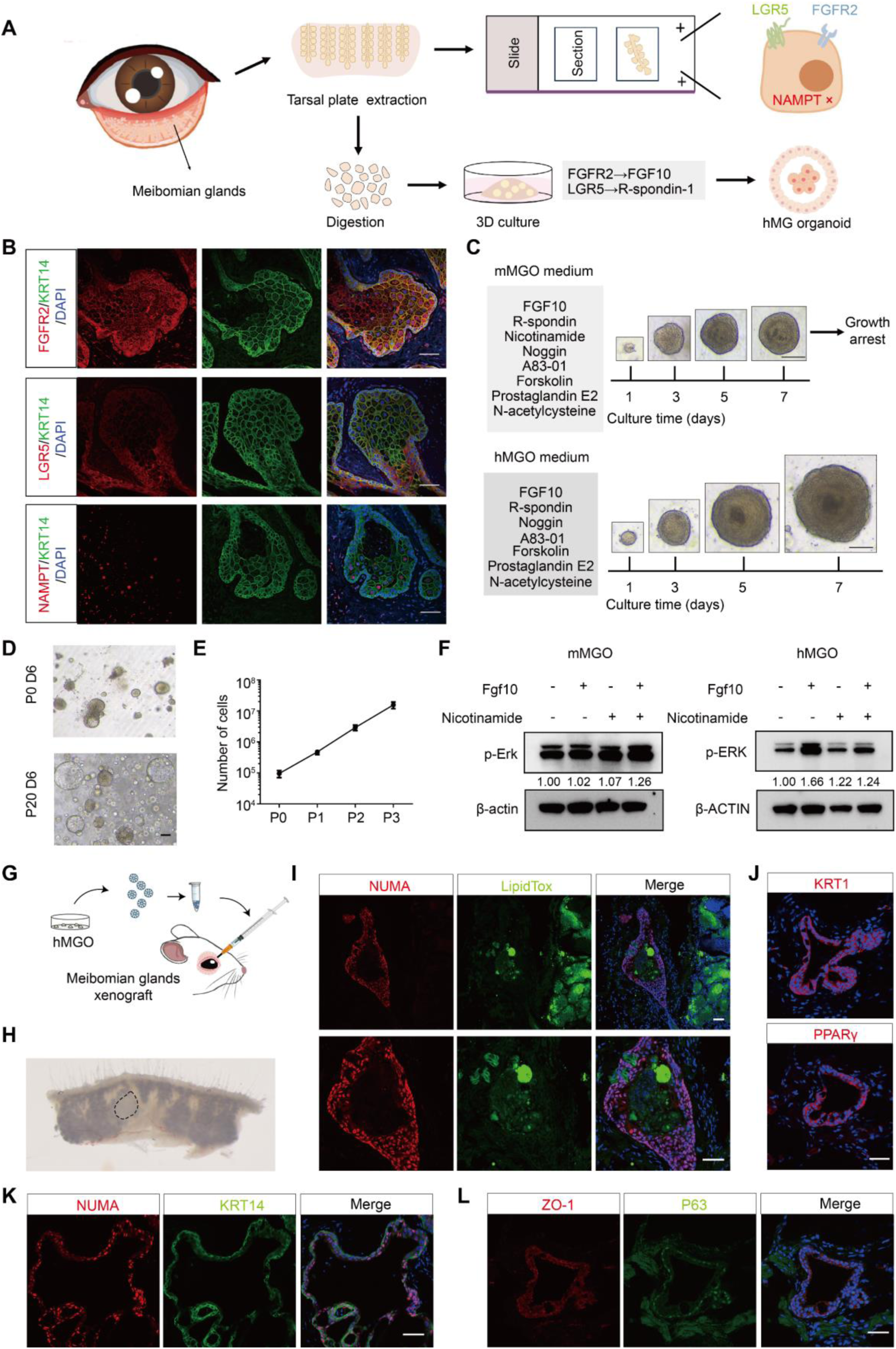
Establishment of human MG organoids for orthotopic transplantation. (A) Schematic diagram of organoid culture from human MG tissue. (B) Immunofluorescent staining for FGFR2, LGR5 and NAMPT in human MG tissue (marked by KRT14). Scale bar, 50 μm. (C) Representative bright-field image of human MG organoids with mMGO medium and hMGO medium at indicated days. Scale bar, 50 μm. (D) Representative images of hMGOs outgrowth from primary organoids to passage 20. Scale bar, 100 μm. (E) The cell number of hMGOs during passage 0-3. (F) Western blotting analysis of phospho-ERK for murine MG organoids and human MG organoids cultured under indicated treatment. β-ACTIN served as a loading control. The relative band intensity that normalized to loading control was marked under each band. n=2 biological replicates. (G) Schematic diagram of human MG organoid orthotopic transplantation in NSG mice. (H) Representative image of hMGOs orthotopic transplantation. Black dashed box indicates region of xenograft. (I) Immunofluorescence staining for human NuMA and LipidTox staining of human engrafted organoids. Lower panel: higher magnifications of upper panel. Scale bar, 50 μm (J-L) Immunofluorescence staining for KRT1, PPARγ (J), human NuMA, human KRT14 (K), ZO-1 and P63 (L) in engrafted MG organoids. Scale bar: 50 μm. See also Figure S5, S6

To further confirm the inhibitory effect of nicotinamide on hMGO growth, we added nicotinamide back to hMGO medium, and found it significantly suppressed cell proliferation and lipid synthesis (Extended Data Fig. 5A-E). Bulk RNA-seq revealed that nicotinamide supplement downregulated cell proliferation and lipid-producing related genes including *PCNA*, *MKI67*, *PPARG* and *SCD* (Extended Data Fig. 5F). GO term and GSEA analysis further confirmed nicotinamide inhibited cell proliferation and PPAR related signaling pathways (Extended Data Fig. 5G-H). Unexpectedly, nicotinamide also downregulated FGF10-FGFR2 signaling (Extended Data Fig. 5H), implying the crosstalk between nicotinamide and FGF10 signaling. To further explore the mechanism, we investigated the phosphorylated ERK (pERK) level in organoids upon FGF10 and nicotinamide treatment. Different with murine MGO, whose pErk level was not altered by nicotinamide, the human MGO’s pERK was quite sensitive to nicotinamide and FGF10: FGF10 treatment significantly upregulated pERK, which was suppressed by nicotinamide subsequently (Fig. 4F). To ensure the necessity of other gradients in hMGO medium, we subtracted each component and evaluated the organoid growth, and confirmed the current hMGO medium is the minimal formula (Extended Data Fig. 5I-K and Extended Data Fig. 6A-B). The hMGO cultured with this medium also preserved the key makers (KRTs and PPARγ) and lipid synthesis capacity, compared with parental tissues (Extended Data Fig. 6C, D).

Dry eye syndrome caused by MGD is a chronic disease that cannot be completely cured theoretically, so we wondering if the transplanted hMGO could generate functional MGs. We transplanted hMGO into the MG sites of immunodeficient nonobese diabetic (NOD) severe combined immunodeficiency (SCID) gamma (NSG) mice (Fig. 4G). After one month, the recipient MGs were collected for analysis. The injected organoids could be observed in the MG tissue (Fig. 4H). The human-derived organoids were confirmed by the staining of human NuMA and KRT14 in the engraftment organoids (Fig. 4I, K). Through LipidTOX detecting, the secreted lipids were detected in the center of the engrafted organoids (Fig. 4I). Further, a similar level of lipid production was observed in the engrafted organoids and the host MGs, confirming that the engrafted organoids acquired functional maturity *in vivo* (Fig. 4I). Additionally, the engrafted cells retained the expression of functional MG biomarkers, including KRT1, KRT14, PPARγ, and P63α, corresponding with the host tissues (Fig. 4J-L). Taken together, the long-term expandable human MG organoids can be engrafted into mice and exhibit features of functional maturation, including lipid production and lineage-specific marker expression.

### Dissection of single-cell atlas in human MGs and organoids

To dissect the cellular composition and heterogeneity in human MGs and organoids, we performed parallel single-cell RNA sequencing (scRNA-seq). The human MG tissues were digested into single cells and subjected to 10x genomics-based scRNA-seq platform (Fig. 5A). After quality control, a total of 23,323 cells were obtained for further analysis and six cell types were identified (Extended Data Fig. 7A, B). Focusing on the epithelial homeostasis, we selected *KRT14*^+^ MG epithelial cells (n=13,684) and further identified nine subclusters according to the established cell markers (Fig. 5B, Extended Data Fig. 7C-G). Based on the high expression of the mature lipid-synthetic marker genes, we identified two types of differentiated acinar cells. Type 1 acinar cells (*FAR2^+^*) are featured by lipid metabolic genes, including *PPARG*, *SCD*, *FAR2* and *MSMO1*. While Type 2 acinar cells (*LYZ^+^*) are featured by secretive genes, including *LYZ*, *SCGB3A1*, *PLA2GA2* and *PIP*. Thus, those two types of acinar cells were identified as mature acinar cells (Fig. 5C, Extended Data Fig. 7G). We also identified other cell lineages of MGs, including MG basal cells (*KRT15*^+^/*SOCS3*^+^; *ID4*^+^/*CCN4*^+^; *UBE2C*^+^/*TOP2A*^+^), transitional cells (*CREB5*^+^/*LAMA4*^+^; *MMP10*^+^/*FST*^+^), and ductal cells (*PIGR*^+^/*OPRPN*^+^; *KRT6A*^+^/*SBSN*^+^; Extended Data Fig. 7D-F).

**Figure 5.**
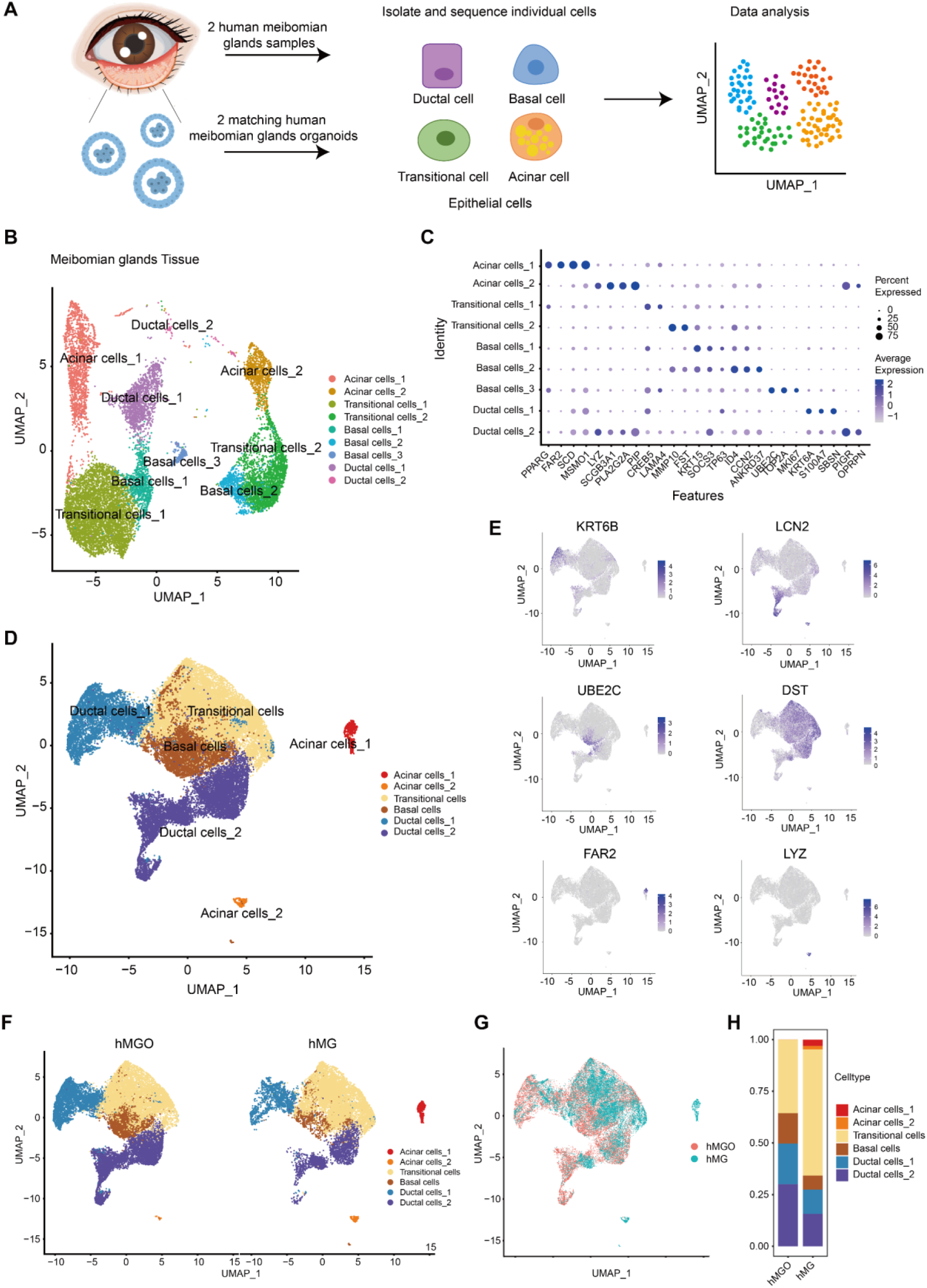
Mapping the single-cell atlas for human MG tissue and organoids. (A) The schematic diagram of single-cell mRNA sequencing for 2 human MG tissue samples and matching organoids. (B) UMAP representation of 9 clusters of human MG epithelial cells (n=12,707). (C) The expression level of MG cell markers and cluster-specific genes in different clusters of human MG tissue. (D) Cell clusters were visualized for human MG organoids and tissue cells. (E) The expression of cluster-specific genes in organoids and tissue cells. (F) UMAP plot showing the parallel cell clusters were identified in organoid and tissue cells. (G) UMAP of human MG cells from organoids (tomato dots) and tissue (cyan dots). (H) The proportion of diverse cell types in human MG organoids and tissue. See also Figure S7

In parallel, we also sequenced 22,077 cells in MG organoids. After aligning with human MG tissues (*KRT14*^+^), the total cells could be aligned and regrouped into five different clusters, namely acinar cells (type1 and 2), ductal cells (type1 and 2), transitional cells and basal cells (Fig. 5D-G). Notably, the type1 acinar cells (*FAR2*^+^) were absent in the organoids, while another type acinar cells (*LYZ*^+^) had a much lower abundance in the organoids (Fig. 5E-G). This indicated that human MG organoids retained relative immature status. Besides, the cell component proportion analysis also revealed an increase in the ductal cells and basal cells in organoids and a decrease in transitional cells (Fig. 5H), indicating human MG organoids preserved more stemness related cells than tissue. Collectively, the conducted single cell atlas described parallel heterogenous cell types between the organoids and MGs, and pinpointed the maturation gap between organoid and tissue.

### Human MG organoids generated featured lipidome

Since we have proved that human MG organoids could reproduce the capacity of lipid synthesis (Extended Data Fig. 6D), we wonder the lipid contents of organoids would resemble those of tissues. In order to depict a broader landscape of lipidome, we initially performed non-targeting lipidomic analysis: using human sebaceous glands (SGs) as control, the lipid samples of human MGs and organoids were extracted and coupled with an electrospray ionization high-resolution mass spectrum (MS) analysis. Then, the normalized intensity profiles of the identified lipid compounds were generated. On Principle component (PCA) score plot, MGs and organoids are distinctly separated from the SGs sample (Fig. 6A), and hierarchical clustering grouped MG tissue and organoids together, confirming organoids shared the similar lipidome with MG tissue (Fig. 6B). Then, we probed the content of subclass lipids in human MG organoids, normal MG tissue, and SG tissue. Distinguished with SG, MG tissue and organoids shared similar level of sphingomyelins (SM), phosphatidylglycerols (PG) and phosphatidylserines (PS). While diacylglycerols (DAG), the lipid content mainly secreted in SG, were much lower in MG tissue and organoids. (Figure 6C, Extended Data Fig. 8A, B). Apart from the similarities, we also noticed the 240 differential lipids between hMG and hMGO (Extended Data Fig. 8C). Among those, FAHFA and TAG, which were detected in human meibum ^28, 29^, were downregulated in hMGO (Extended Data Fig. 8D).

**Figure 6.**
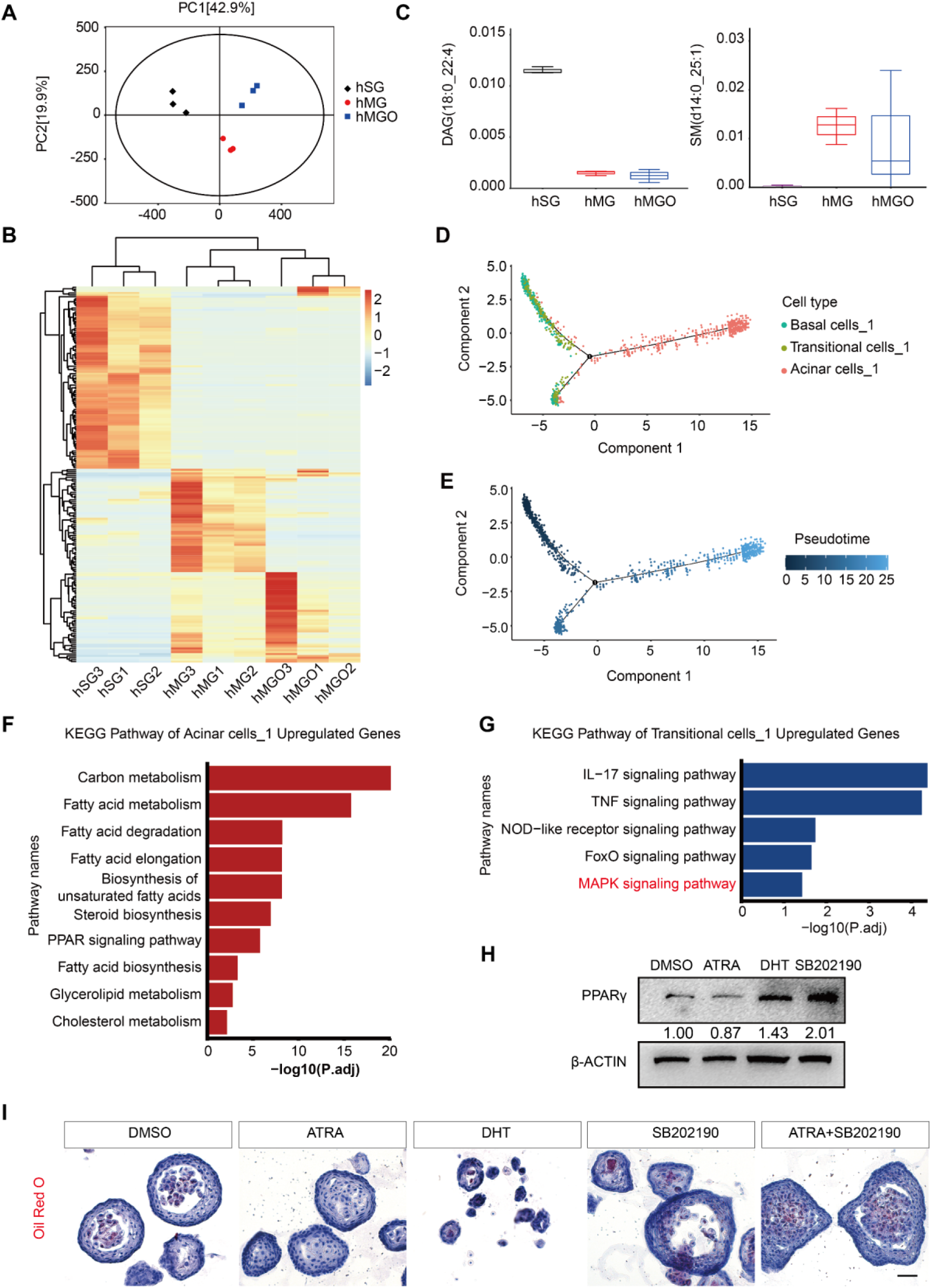
MAPK signaling inhibition promotes the maturation of human MG organoids. (A) PCA plot of human sebaceous gland (hSG), meibomian gland (hMG) and meibomian gland organoids (hMGO) in lipidomics analysis. 3 independent donors in each group. (B) Heatmap of differential lipid species among hSG, hMG and hMGO. (C) Relative value of DAG (18:0_22:4) and SM(d14:0_25:1) in hSG, hMG and hMGO. (D) Position of the MG cells in different cell clusters along pseudotemporal trajectory. (E) Ordering of MG tissue cells along a pseudotemporal trajectory. MG cells are colored by pseudotime (F-G) KEGG analysis of upregulated genes (F) and downregulated genes (G) that differentially expressed in MG acinar cells, compared with MG transitional cells. (H) Western blotting analysis of PPARγ for human MG organoids cultured under indicated treatments: 1% DMSO, 100μ ATRA, 10μM DHT, and 0.5μM SB202190. β-ACTIN served as a loading control. The relative band intensity that normalized to loading control was marked under each band. n=2 biological replicates. (I) The Oil Red O staining of human MG organoids cultured under the treatment of ATRA, DHT and SB202190. Scale bar: 50 μm. See also Figure S8

In summary, consistent with scRNA-seq data above, the lipidomic comparison between MG tissue and organoids not only displayed the overall consistency, but also suggested the maturation gaps, such as terminal differentiated cell types and featured lipids.

### MAPK signaling inhibition boosted human MG organoids maturation

Since the hMGO are less mature than MG tissues, we reasoned that the underlying signals in MG tissues could enhance organoid maturation. To unveil the mechanism of acinar cell maturation, lineage trajectory analysis was performed based on the scRNA-seq data of MG tissues. On the pseudotime plot, basal cells, transitional cells and acinar cells of MGs distributed along the pseudotime trajectory. The type1 acinar cells (*FAR2*^+^), which were identified as the mature MG cells, was located at the rightmost of the trajectory (Fig. 6D, E), suggesting that the differentiation path of basal cells-transitional cells-acinar cells. Then we analyzed the signaling pathway enriched in acinar and transitional cells, in order to find key signaling pathways guiding acinar cell maturation. In acinar cells, multiple lipid synthesis-related signaling pathways were activated (Fig. 6F), while the main signaling pathways enriched in transitional cells were inflammation related pathways such as the IL-17, TNF, NOD-like receptor (NLR), FoxO, and MAPK signaling pathway (Fig. 6G). Interestingly, MGD patients were generally associated with increased tear cytokines, such as TNF-α, IL-17A, and IL-1β ^30^. Reducing the inflammatory cytokines could relieve both the signs and the symptoms of MGD ^30, 31^. Thus, we exposed human MG organoids to the inhibitors of the IL-17, TNF, NLR, FOX, and MAPK signaling pathways and attempted to enhance the lipid production capacity of human MG organoids. Notably, a selective p38MAPK signaling pathway inhibitor, SB202190, showed the potential to facilitate MG organoid maturation.

As previously reported, the p38MAPK signaling pathway contributes to cell apoptosis and the autophagy of adipogenic cells and plays a crucial role in inflammation-induced MGD ^32, 33^. Treatment with SB202190 could significantly upregulate the expression of PPARγ in organoids, and it is more effective than DHT in this regard (Fig. 6H). Using Oil Red O staining, we observed that upon SB202190 stimulation, the produced lipids, which were originally in the cytoplasm, were released into the cavity of the organoids (Fig. 6I). Although the ATRA treatment decreased lipid secretion, SB202190 mitigated the inhibiting effect of ATRA (Fig. 6I). These results confirmed that SB202190, a selective inhibitor of the p38MAPK signaling pathway, could induce the maturation of MG organoids.

The mature meibum is featured by ultra-long chain lipids with fatty acid and/or fatty alcohol chains up to C34-C36, including cholesteryl esters (CE), (O)-Acylated u-hydroxy fatty acids (OAHFA) and wax esters (WE)^34, 35^. However, the non-targeting lipidomic analysis could not robustly detected those major components, especially for the WE. To depict the maturity of human meibomian organoids, targeting lipidomic analysis was performed to evaluate the content of key lipids of meibum, including CE, OAHFA and WE. Overall, human MG organoids (hMGO) resembled the lipidomic features of human MG tissue (hMG) in PCA plot, compared with human periocular skin tissue (hST) (Extended Data Fig. 9A). Then we used heatmaps to delineate the similarity between hMG and hMGO on CE, OAHFA and WE. Compared with negative control hST, the average content of those featured lipids in hMG and hMGO is higher than hST (Extended Data Fig. 9B-D). However, we also observed the variant lipidomic performance of hMGO, which showed less mature phenotype (such as hMGO1), suggesting the improving space for hMGO culture system.

## Discussion

Efforts have been made to establish 3D meibomian culture model, including slice culture model and explant culture methods ^36–40^. Although these models preserved the tissue architectures and functions, such as lipid production, they could not sustain for more than 7 days *in vitro*. In this study, we achieved long-term expansion of murine and human MG organoids, which retained all cell lineages and capacity of lipid synthesis. Thus, the MG organoids enable precise drug response and functional engraftment. In addition, MG organoids also assist to untangle the underlying mechanism of tissues homeostasis and dysfunction. We found that FGF10 signaling is crucial for organoid expansion and MGs regeneration; NAMPT was barely expressed in human MG tissue, and nicotinamide specifically hindered human MG organoid expansion, indicating the distinct NAD metabolism between mouse and human; Single-cell atlas of human MG epithelial cells indicated MAPK signaling inhibition is required for acinar cell differentiation, thus treated with p38MAPK signaling pathway inhibitor, SB202190, promote organoid maturation.

MGD is the leading cause of dry eye disease. Clinically, the main treatments for MGD include cleaning the eyelids through hot compresses and massages, instilling artificial tears containing lipid components, and using antibiotic drugs to reduce inflammation. Currently, the development of artificial tears is prevailing. In order to make the artificial tears as similar as possible to the composition of tears, various substances are added, including mineral oil and phospholipids^41^. We identified that FGF10 supported both murine and human MG organoid expansion. And FGF10 eye drops also rescued the ATRA induced MGD. (Fig. 1 and Fig. 2). Our data suggested that the FGF10 signaling is not only required for MG develop and homeostasis ^19, 42, 43^, but also prevent the progress of MGD. Therefore, FGF10 eye drops is the promising reagent for human dry eye disease, which need more human relevant tests for its application. In addition to those physiological restoration, FGF10 treatment also increased the proliferation of acinar cells, instead of ductal cells (Fig. 2N). Consistent with our observation, FGFR2 deletion severely affected proliferation and differentiation of MG acinar cells but affected MG ductal cells to a lesser extent ^43^. We also found that the proliferating cells in acini are mainly located at the edges (Extended Data Fig. 2A), which were likely basal cells. Consistent with this data, the MG organoid also showed proliferative Krt5^+^ basal cells under the FGF10 containing medium (Fig. 1E). Hence, it is conceived that basal cells are the major proliferative cells that responds to FGF10 signals. NAMPT generates nicotinamide mononucleotide from nicotinamide and 5’-phosphoribosyl-1-pyrophosphate, thereby catalyzing the rate-limiting step in the mammalian NAD salvage pathway^18^. Therefore, the bonus of nicotinamide largely depends on the activity of NAMPT. We found the distinct expression level of NAMPT in MGs between mouse and human (Fig. 4B). And the addition of nicotinamide into human organoid culture medium suppressed the organoid growth (Fig. 4C). Previous studies also revealed that high doses of nicotinamide may exert adverse effects, such as DNA damage, increased intracellular ROS, spindle defects, and mitochondria dysfunction ^44–46^. In this study, we found nicotinamide inhibited the activity of FGF10 signaling through decreasing the pERK level in human organoids. But how nicotinamide crosstalk with upstream of FGF10 signaling need further investigation. It was reported that the prevalence of dry eye disease was higher in female than male^47^. Considering the nicotinamide is frequently used in cosmetics, it’s likely that excess nicotinamide exposure is also an emerging risky factor of MGD.

MAPK signaling participants in various cellular programs like proliferation, differentiation, transformation and apoptosis ^48^. Leveraging scRNA-seq analysis, we found that MAPK signaling pathway was significantly enriched in transitional cells (immature acinar cells), compared to mature acinar cells (Fig. 6G). And identified SB202190, a selective p38MAPK inhibitor, promoted lipid production in MG organoids (Fig. 6I). Suggesting MAPK signaling pathway inhibition is required for the differentiation of mature acinar cells. Similar to adipocytes, the p38MAPK signaling was more active in preadipocytes than adipocytes ^49^. Consistently another selective p38MAPK inhibitor, SB203580, could block IL-1β-induced MGD and restore its functions in rat ^50^. These findings indicate that MAPK signaling inhibition enhances the functions of terminal differentiated cells, and could be considered as the potential targets for MGD. However, whether MAPK inhibition affect the self-renewal of basal cells remains further investigation.

MG organoids provide the opportunity of cell therapy for MG atrophy. Given that MG atrophy is the terminal stage of MGD, when MG stem cell could not regenerate the whole MGs, the orthotopic transplantation might be the only choice for those patients. Successfully engrafting MG organoids is the first step for potential regenerative therapies ^51^. The established MG organoid culture and differentiation protocol would provide the “seed cells” for bioengineered MGs grafts. Efforts should be done to improve the orthotopic transplantation efficiency and reduce the immune rejection effects caused by xenotransplantation. In conclusion, we established functional, long-term-cultured MG organoid cultures for MG homeostasis and diseases research, which provides a useful platform for the development of associated medical interventions and regenerative medicines.

In MGs, the acini are connected to the main duct by small ducts, so that the lipid produced by the meibocytes could be transported through the ducts to the eyelid. Although our organoid model contained all cell types of MGs, the ductal cells resided with basal cell to form the out layer of the organoid, losing its function to drain out the meibum. Additional optimization of MG organoid establishment is required to develop ducts-acinus structure.

In addition, we showed that the human MG organoids engrafted upon orthotopic transplantation, and the transplanted organoids could produce comparable lipids to its neighboring tissue. However, whether the MG organoid transplantation could rescue the MG dysfunction remain further investigation.

## Methods

### Ethical regulation compliance

This research complies with all relevant ethical regulations. The animal study was approved by the Laboratory Animal Center, School of Life Sciences, Fudan University and was performed in accordance with the statement of the Association for Research in Vision and Ophthalmology (ARVO) entitled “Use of Animals in Ophthalmic and Vision Research” (EENT-IRB20220301). The research was conducted in accordance with the Declaration of Helsinki, and the use of human tissues for this study was approved by the Ethics Committee of EENT hospital of Fudan University (EENT-IRB20220222).

### Tissue collection and MG organoid cultures

For the murine MG organoids, six to eight-week-old wild-type C57BL/6 mice were used for the culture of the murine MG organoids. The eyelids of mice were obtained as well as the tarsal plates were isolated. All of the skin, muscles, and connective tissues were removed under a dissecting microscope, and the tissues were cut into approximately 1 mm^3^ pieces. Then, these were enzymatically digested in 1 mg/mL collagenase I (Sigma-Aldrich, C9407) with 10 μM Y-27632 (Sigma-Aldrich, Y0503) and 20 U/mL DNase I (Sigma-Aldrich, DN25) for 30 min at 37° C. For the human MG organoids, MGs from cadaveric human aged 30–70 years were obtained from an eye bank. The tarsal plates were digested using 1 mg/mL collagenase I with 10 μM Y-27632 and 20 U/mL DNase I for 90 min at 37° C, and the MGs were squeezed out using MG forceps. The digestion solution was washed twice with Advanced DMEM/F12 (Gibco), and the cells were placed in a 30 μL Growth Factor Reduced Basement Membrane (Corning, 356231) in a 24-well plate.

The mouse expansion medium contained Advanced DMEM/F12 (Gibco, 12634028) supplemented with 1xB27 (Gibco, 17504044), Glutamax (Gibco, A1286001), HEPES (Gibco, 15630130), 100 U/mL Penicillin Streptomycin (Gibco, 15070063), 1.25 Mm N-acetylcysteine (Sigma-Aldrich, USA), 10 mM nicotinamide (Sigma-Aldrich, 616911), 0.5 μM A83-01 (OrganRegen, C909910), 1 μM PGE2 (OrganRegen, C323646), 1 μM FSK (OrganRegen, C665752), 100 ng/mL FGF10 (OrganRegen, 816-FGF), 100 ng/mL Noggin (OrganRegen, 807-NOG), and 500 ng/mL Rspo-1 (OrganRegen, 861-RS1). The nicotinamide was removed from the murine medium, which constituted the human MG organoid medium. After six days of seeding, the organoids were mechanically dissociated into small fragments using Pasteur pipettes and digested using TrypLE Express with 1 mM EDTA. The dissociated cells were then embedded in fresh Matrigel; next, the murine organoids were passaged at a rate of 1:3–1:4, while human organoids were passaged at a rate of 1:4–1:5. The organoids were incubated at 37° C with 5% CO2, and the medium was changed every two to three days.

For the drug testing, the MG organoids were cultured for three days and incubated with 0.1% DMSO (Sigma-Aldrich, D5879), 100 μM ATRA (Sigma-Aldrich, R2625), 100 μM 13-cis RA (Sigma-Aldrich, R3255), 10 μM AZM (MedChem Express, HY-17506), 10 μM Rapamycin (MedChem Express, AY-22989), 10 μM DHT (Sigma-Aldrich, D-073), and 0.5 μM SB202190 (OrganRegen, C152121) for three days.

### Colony formation assay

The organoids were dissociated into single cells, and a total of 2,000 cells of organoids were seeded per 96-well plate and cultured for six days. The number of organoids was counted, and the colony-forming efficiency was calculated by the following formula: (number of organoids)/2000 × 100%.

### Immunofluorescence staining

The organoids were harvested, fixed with 4% paraformaldehyde for 30 min at room temperature, and the MGs were isolated and fixed overnight at 4° C. For the frozen section, samples were dehydrated in 30% sucrose and embedded in optimum cutting temperature compound (SAKURA, 4583). For the paraffin section, samples were dehydrated using ethanol of increasing gradients, washed in xylene, and embedded in paraffin. The embedded samples were cut before staining. Hematoxylin and eosin (H&E) staining was performed for the organoids and tissues according to the manufacturer’s instructions.

Briefly, for immunofluorescence staining of the organoids and tissues, after washing with PBS, the samples were incubated at 4°C overnight in 0.3% Tritonx-100 and 5% BSA with primary antibodies. After being washed with PBS thrice, the organoids were incubated with secondary antibodies at room temperature for one hour. The nuclei were labelled using Hoechst 33258 (1:2000, Invitrogen, USA), and images were acquired with a confocal microscope (OLYMPUS, FV3000). Information about the antibodies is provided in Supplementary table 1.

### Oil Red O staining

An Oil Red O staining kit (Beyotime, C0175S) was used to assess the lipid production of the organoids and tissues according to the manufacturer’s protocol. The frozen sections of the organoids and tissues were stained with a fresh Oil Red O solution for 12 min before being washed twice by 60% isopropanol for 3–5 s. After rinsing with PBS for 5 min, hematoxylin was used for staining the cell nuclei, and images were captured by the cellSens system of the Olympus microscope.

### LipidTOX staining

After being washed in PBS for 5 min, the frozen sections of the MGs were incubated with the LipidTOX neutral lipid satin (1:1000, Thermofisher, H34475) in 5% BSA at room temperature for at least 30 min. Then, images were acquired and evaluated using a confocal microscope (OLYMPUS, FV3000).

### Real-time Quantitative PCR (qPCR) analysis

The organoids of the 24-well plate were collected, and the total RNA was extracted using the TRIzol reagent (Invitrogen, USA) according to the manufacturer’s instructions. The PrimeScript™ RT Reagent Kit with gDNA Eraser (TaKaRa, Japan) was used for the reverse transcription of cDNA. The qPCR analysis was performed using a QIAantiNova SYBR Green PCR Kit (Qiagen, Germany) with a protocol of two minutes of initial activation at 95°C, 40 cycles of 5 s of denaturation at 95°C, and 20 s of annealing and extension at 60° C. The mRNA expression levels were determined using the 2^-ΔΔCt^ method, in which the target gene was normalized to β-actin to calculate the relative mRNA fold change. The sequences of the target-gene-specific primers are presented in Supplementary table 2.

### Western blotting

The organoids were harvested and washed with PBS, and the total protein was isolated using the RIPA buffer (Beyotime Biotechnology, China). The protein concentration was measured using a BCA protein assay kit (Beyotime Biotechnology, China). The protein was separated using 4–20% SDS-polyacrylamide gel electrophoresis (GenScript, China) and transferred to a nitrocellulose membrane (Millipore, USA). The membranes were blocked with 5% nonfat powdered milk and incubated with primary antibodies at 4°C overnight. After thorough washing with Tris-buffered saline with Tween (TBST), HRP-conjugated secondary antibodies were added for incubation with the membranes at room temperature for one hour, using β-actin as an internal reference. The protein expression levels were semi-quantified with ImageJ software (NIH, Washington, DC). Information about the antibodies is presented in Table S2.

### mRNA sequencing

#### Tissue isolation and sample preparation

The total RNA of the organoids was collected and extracted as described above. For tissues, the MGs and lacrimal glands were isolated from wild-type mice and digested in 100 μg/mL Liberase TM (Sigma-Aldrich, 5401020001) and 100 U/mL DNase I for one hour at 37°C. After incubating the sample with 0.25% Trypsin for five minutes, single-cell solutions were collected and filtered before sorting. The EPCAM-FITC antibodies (1:200, Biolegend, 118208) were stained to isolate the epithelial cells of the MGs and lacrimal glands, and the isolation was performed using a BD FACSAria™ Fusion sorter. The EPCAM^+^ cells were collected, and their total RNA was extracted as described above. For murine and human organoids, established organoid lines (>passage 5) cultured for 7 days were collected, and the total RNA was extracted as described above.

#### mRNA sequencing and data analysis

After RNA quality control and sequencing, a total of 1 μg of RNA was used to prepare RNA-sequencing libraries using an Illumina HiSeq platform, and more than 40 million readings were obtained for each sample. FastQC (https://www.bioinformatics.babraham.ac.uk/projects/fastqc/) was used to check the quality of the raw sequencing data, and HISAT2 (https://daehwankimlab.github.io/hisat2/) was used to match the readings to the mouse genomes with default parameters. Further, BAM files were sorted using SAMtools, while count matrices were generated using HTseq v 0.11.2.

The downstream analysis was carried out in R v 4.1.3. DESeq2 v 1.34.0 was used to identify differentially expressed genes (DEGs), and genes with a p-value of 0.01 or lower were regarded as DEGs. Finally, clusterProfiler v 4.2.2 was used to perform GSEA, and gene expression heatmaps were generated using pheatmap v 1.0.12

### Single-cell mRNA sequencing

#### Single-cell RNA sequencing and data preprocessing

To prepare single-cell suspensions, two human MG tissue samples (a 68-year-old male and a 62-year-old male donors) were isolated as described above, as well as two organoid samples were dissociated using TrypLE for 6 min at 37°C. Single-cell sequencing was performed using the 10x Genomics platform. The libraries were input into the Illumina Novaseq 6000 platform for PE150 sequencing.

The sequencing data were first processed through the official 10x Genomics pipeline Cell Ranger v 6.0.2 and then matched to the hg38 human reference genome. Raw gene expression matrices were loaded into R using Seurat v 4.3.0. Poor-quality cells with abnormally high unique-feature counts or mitochondrial gene expression were removed. For the remaining cells, doublet detection and filtering were performed using DoubletFinder. After quality control, 15,635 cells from the tissue samples and 19,202 cells from the organoid samples were used for subsequent analysis.

#### Unsupervised clustering analysis

For clustering analysis, cells from different samples were merged and normalized using the Seurat function NormalizeData. Next, FindVariableFeatures was used to identify highly variable genes, and the principal components were estimated after scaling. The authors used FindNeighbors with the first 20 components to allocate cells, and FindClusters was employed to cluster the cells. UMAP dimensionality reduction was then performed using RunUMAP. The cell types were annotated based on the expression pattern of the DEGs and the well-known cellular markers from the literature.

#### Data integration

The Harmony package was utilized to integrate the data from two tissue samples, followed by LIGER documentation to integrate the single-cell data from different tissue and organoid samples.

#### Differential expression analysis and pathway enrichment analysis

To identify the DEGs of different cell types, the Wilcox algorithm was used for statistical testing via the Seurat functions FindAllMarkers and FindMarkers. KEGG enrichment analysis was carried out using the clusterProfiler R package.

### Non-targeting lipidomic analysis

#### Lipid extraction

The hMG, hMG organoids and hSG were used for lipid extraction. For hMG, 8.8 mg freeze-dried meibum were collected by squeezing hMGs from 3 healthy male donors, and 200 μL water and 480μL extract solution (MTBE:MeOH=5:1) was added for extracting. For hSG, facial sebum was wiped and collected by swab from another three healthy male donors, 500 μL water and 1200 μL extract solution (MTBE:MeOH=5:1) was suppled sequentially. Then, hMG and hSG samples were homogenized at 35 Hz for 4 min and sonicated for 5 min. The homogenization and sonication cycles were repeated for 3 times. For hMG organoids, 200 μL water was added into the MG organoids of passage 5. After 30s vortex, the sample were freezed with liquid nitrogen for 1 min and thawed. After this cycle repeated for 3 times, organoids were sonicated for 10 min, and 480 μL extract solution (MTBE:MeOH=5:1) was performed. hMG, hSG and organoids were incubated at −40°C for 1 hour, and the supernatant was transferred and dried in a vacuum concentrator at 37°C. Finally, the solution (DCM: MEOH=1:1, containing internal standard) was suppled into dried samples for reconstituted. The supernatant was used for LC/MS analysis.

#### LC-MS/MS analysis

LC-MS/MS analysis used UHPLC system (Vanquish, Thermo Fisher Scientific, USA). The mobile phase A consisted of 40% water, 60% acetonitrile, and 10mmol/L ammonium formate, and the mobile phase B consisted of 10% acetonitrile and 90% isopropanol, which was added with 50mL 10mmol/L ammonium formate for every 1000ml mixed solvent. Elution gradient was used for analyzing: 0∼1.0 min, 40% B; 1.0∼12.0 min, 40%∼100% B; 12.0∼13.5 min, 100% B; 13.5∼13.7 min, 100%∼40% B; 13.7∼18.0 min, 40% B.

The QE mass spectrometer was used to acquire MS/MS spectra on data-dependent acquisition mode in the control of the acquisition software (Xcalibur 4.0.27, Thermo Fisher Scientific, USA). And the ESI source condition were set respectively: sheath gas flow rate as 30 Arb, Aux gas flow rate as 10 Arb, capillary temperature 320°C (positive), 300°C (negative), full MS resolution as 70000, MS/MS resolution as 17500, collision energy as 15/30/45 in NCE mode, spray Voltage as 5 kV (positive) or −4.5 kV (negative).

#### Data preprocessing and annotation

The CentWave algorithm in XCMS was used for peak detection, extraction, alignment, and integration. Lipid identification was achieved through a spectral match using LipidBlast library, which was developed using R and based on XCMS.

### Targeting lipidomic analysis

#### Lipid extraction

Lipids were extracted from frozen tissues using a modified version of the Bligh and Dyer’s method as described previously^52^. Briefly, tissues were homogenized in 900 µL of chloroform: methanol: MilliQ H2O (3:6:1) (v/v/v). The homogenate was then incubated at 1500 rpm for 30min at 4℃. At the end of the incubation, 350 µL of deionized water and 300 µL of chloroform were added to induce phase separation. The samples were then centrifuged and the lower organic phase containing lipids was extracted into a clean tube. Lipid extraction was repeated once by adding 500 µL of chloroform to the remaining aqueous phase, and the lipid extracts were pooled into a single tube and dried in the SpeedVac under OH mode. Samples were stored at −80℃ until further analysis.

#### Lipidomic analyses

Lipidomic analyses were conducted at LipidALL Technologies using a Shimadzu Nexera 20-AD HPLC coupled with Sciex QTRAP 6500 PLUS as reported previously^53^. Separation of individual lipid classes of polar lipids by normal phase (NP)-HPLC was carried out using a TUP-HB silica column (i.d. 150x2.1 mm, 3 µm) with the following conditions: mobile phase A (chloroform: methanol: ammonium hydroxide, 89.5:10:0.5) and mobile phase B (chloroform: methanol: ammonium hydroxide: water, 55:39:0.5:5.5). MRM transitions were set up for comparative analysis of various polar lipids. Individual lipid species were quantified by referencing to spiked internal standards. d9-PC32:0(16:0/16:0), d9-PC36:1p(18:0p/18:1), d7-PE33:1(15:0/18:1), d9-PE36:1p(18:0p/18:1), d31-PS(d31-16:0/18:1), d7-PA33:1(15:0/18:1), d7-PG33:1(15:0/18:1), d7-PI33:1(15:0/18:1), d7-LPE18:1, C17-LPI, C17-LPA, C17-LPS, C17-LPG, C14-BMP, d7-CS were obtained from Avanti Polar Lipids. GM3-d18:1/18:0-d3 was purchased from Matreya LLC. Free fatty acids and OAHFA were quantitated using d31-16:0 (Sigma-Aldrich) and d8-20:4 (Cayman Chemicals).

Glycerol lipids including diacylglycerols (DAG), triacylglycerols (TAG), cholesteryl esters (CE), wax esters (WE) were quantified using a modified version of reverse phase HPLC/MRM^54^. Separation of neutral lipids were achieved on a Phenomenex Kinetex-C18 column (i.d. 4.6x100 mm, 2.6 µm) using an isocratic mobile phase containing chloroform:methanol:0.1 M ammonium acetate 100:100:4 (v/v/v) at a flow rate of 300 µL for 10 min. Levels of short-, medium-, and long-chain TAGs were calculated by referencing to spiked internal standards of TAG(14:0)3-d5,TAG(16:0)3-d5 obtained from CDN isotopes, respectively. DAGs were quantified using d5-DAG17:0/17:0 as internal standards (Avanti Polar Lipids), CE and WE were calculated by referencing to spiked internal standards of d6-C18:0 cholesteryl ester (CE), WE44:1(13C18:1/26:0) obtained from CDN isotopes, respectively.

### ATRA-induced MGD animal model

#### Model establishment and eye drop administration

Six-to eight weeks-old male wild-type C57BL/6 mice, weighing about 20 g each, were used for creating the MGD animal model. They were treated with 10 mg/kg/day of ATRA through gavage for 30 consecutive days to damage their MGs. Next, PBS containing a 10 ng/mL FGF10 solution was topically applied to their right eyes three times a day (at 10:00, 15:00, and 20:00; 5 μL of the solution was administered each time) from Day 0 (n > 3, Figure 2). The left eye was treated with PBS as a control. Additionally, PBS-treated wild-type C57BL/6 mice were used as negative controls (n > 3).

#### Ophthalmologic examinations of the MGD mice

The damage caused by MGD to the corneal epithelium was evaluated through the corneal fluorescein staining (CFS) score, which was performed by dripping 5 μL of sodium fluorescein (0.5%) on the cornea, and excess lipids were wiped away. Images were captured using a slit lamp microscope after three blinks in the cobalt blue channel. The extent of the staining was scored on a scale of 0–3 for the five areas of the cornea (upper, lower, nasal, temporal, and optical diameters), and the total score for each mouse were recorded and analyzed.

Next, a phenol red thread test was performed to measure tear secretion ability. After 20 s of placing the phenol red test in the lower eyelid of the mouse eye, the length of the section whose color changed from yellow to red was measured with a ruler, reflecting tear secretion ability.

To analyze the changes in MG length in different groups, the murine tarsal plates were isolated, and their images were captured. The longest axis of individual MGs was measured using the Image J software, and the average length of the glands in each tarsal plate was calculated.

### Organoid transplantation

For organoid preparation, a 24-well plate of organoids was harvested and resuspended in a 20 μL mixture consisting of half medium and half Matrigel. The organoids were kept on ice until transplantation.

The mouse xenograft model was established using six-week-old NSG (NOD-Prkdcscid Il2rgem1/Smoc) mice obtained from Shanghai Model Organisms. They were kept on a 12 h light/12 h dark cycle before and throughout the study. After being anesthetized with 0.2 mL/10g Avertin through intraperitoneal injection, their upper and lower MGs were exposed, and 5 μL of the cold organoids were injected into the MG tissue using an insulin syringe. After a month of observation, the mice were euthanized, and their MGs were extracted for histological examination.

### Statistical analysis

Three iterations of each experiment were performed, and the statistical analysis was carried out using the GraphPad Prism software (La Jolla, CA, Version 9). The data were shown as mean ± SEM, and the Student’s *t*-test was used for the comparison between the two groups. *p*-values of less than 0.05 were considered statistically significant.

## Data availability

- Single-cell and bulk RNA-seq data have been deposited at NCBl SRA database and are publicly available with the accession number: SRP497138. Lipidomics raw data have been deposited to the MetaboLights with the accession number: MTBLS9799, and can be fast downloaded from Google Drive with the link: https://drive.google.com/drive/folders/1mb84LM6ZEaX-naEBJ24hQx9nLBJ7K7Ix?usp=sharing.
- This paper does not report original code.
- Any additional information required to reanalyze the data reported is available from the lead contact upon request.

## Acknowledgements

The authors thank Jun Xiang and Jiayu Gu for their support with donor tissue acquisition. This study is sponsored by bioGenous BIOTECH. Jiaxu Hong was supported by the National Science Fund for Distinguished Young Scholars (82425015), the National Natural Science Foundation of China (82171102), the National Key Research and Development Program of China (2023YFA0915003), the Shanghai Medical Innovation Research Program (22Y21900900), the “Dawn” Program of Shanghai Municipal Education Commission China (24SG11), the Shanghai Science and Technology Innovation Action Plan for Cell and Gene Therapy (24J22800500), and the Shanghai Science and Technology Innovation Action Plan for Advanced Materials (24CL2900802).

## Author contributions

C.Y., X.W., J.W., J.H. and B.Z. conceived the study; C.Y., X.W., J.W., X.W., Z.X., X.W., Y.M., Y.Z., X.Z., Q.L., J.X., C.Z., X.S. and X.Z. performed the experiments; J.H. and B.Z. supervised the work; and C.Y., X.W., J.W., J.H. and B.Z. wrote the manuscript.

