## Supplementary figure1-9 for "Human meibomian gland organoids to study epithelial homeostasis and dysfunction"

### Extended Data Fig. 1

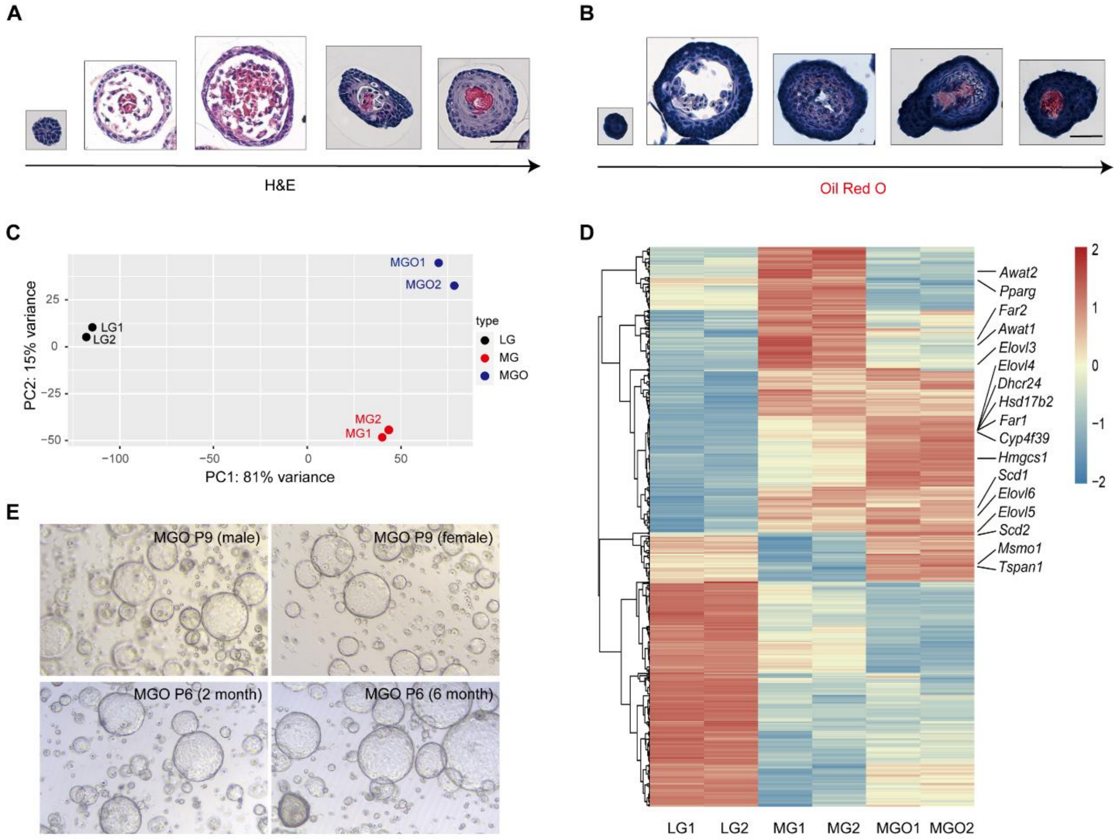


**Extended Data Fig. 1 | Establishment of a long-term 3D murine MG organoid**

(A) Representative H&E staining of single organoid from early to later stage of lifespan. Scale bar, 50 μm.

(B) Representative Oil Red O staining images of single organoid from early to later stage of lifespan. Scale bar, 50 μm.

(C) PCA plot of lacrimal gland (LG), meibomian gland (MG) and meibomian gland organoids (MGO) in bulk RNA-sequencing.

(D) Global transcriptomic analysis of LG, MG and MGO. Key genes of meibum biosynthesis have been indicated.

(E) Bright field image of mouse MGO isolated from different sex and age background.

### Extended Data Fig. 2

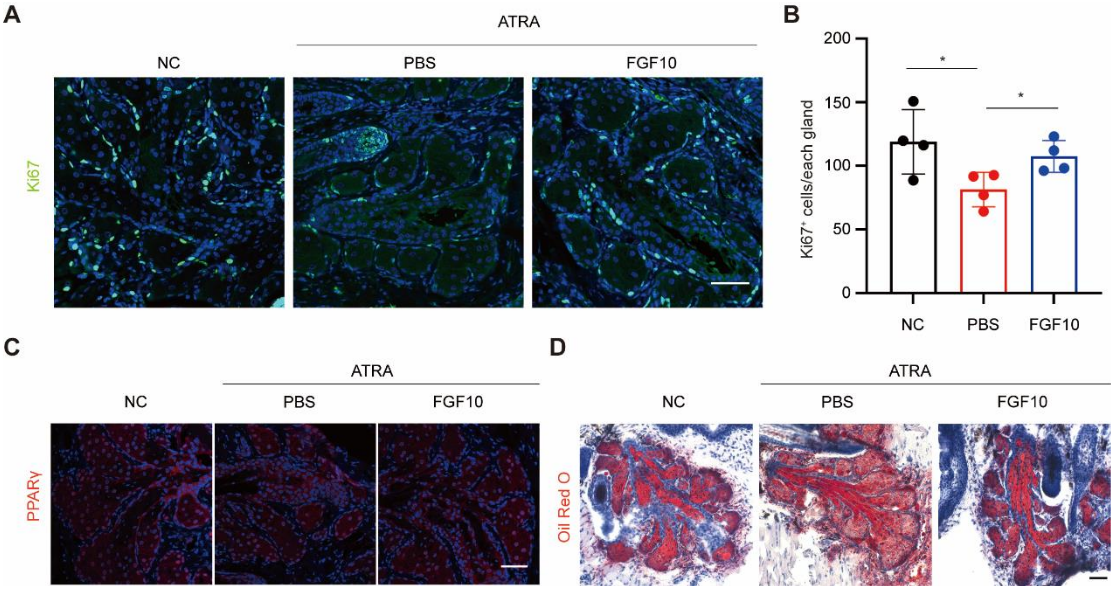


**Extended Data Fig. 2 | FGF10 alleviated the development of MGD**

(A) Immunofluorescent staining for the expression of Ki67 in indicated groups. Scale bar, 50 μm.

(B) Quantification of the Ki67$^{+}$cells in each meibomian gland of indicated groups. Data were represented as mean ± SEM. n=4. Unpaired two-tailed Student's *t* test: * $p$<0.05. Scale bar, 50 μm.

(C) Immunofluorescent staining for the expression of PPARγ in indicated groups. Scale bar, 50 μm.

(D) Oil Red O staining of the MGs tissue in indicated groups. Scale bar, 50 μm.

#### Extended Data Fig. 3

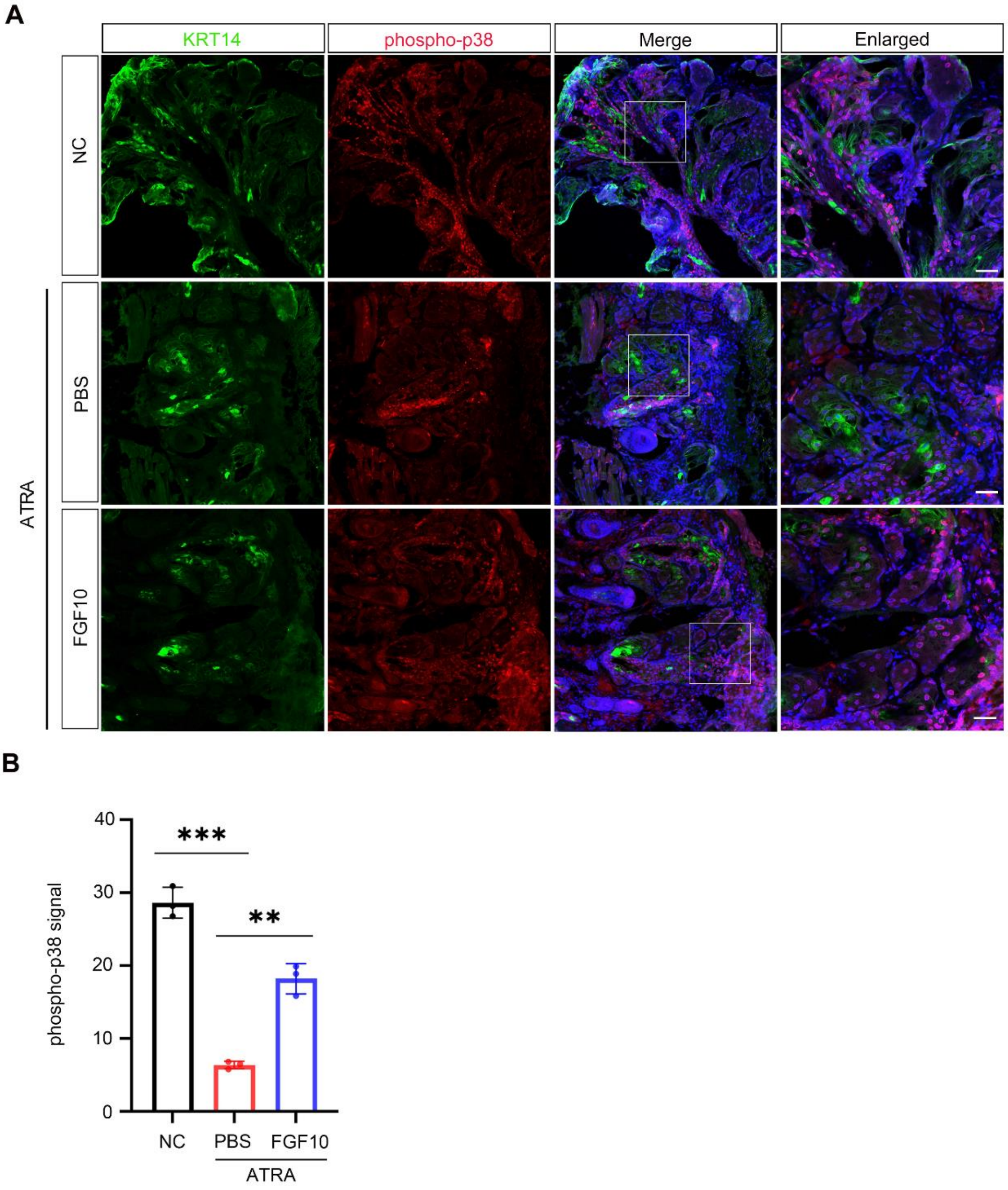


**Extended Data Fig. 3 | FGF10 restore the ATRA-inhibited FGF10-p38 signaling.**

(A) Immunofluorescent staining for KRT14 (Meibomian epithelial cells marker) and phospho-p38 (Activated p38-MAPK signaling pathway marker) of indicated groups. Scale bar, 25μm.

(B) Quantification of the mean fluorescence intensity of phospho-p38 in meibomian gland epithelial cells in indicated groups. Data were represented as mean ± SEM. n=3 for each group. Unpaired two-tailed Student's t test: **p<0.01,

*** p<0.001.

#### Extended Data Fig. 4

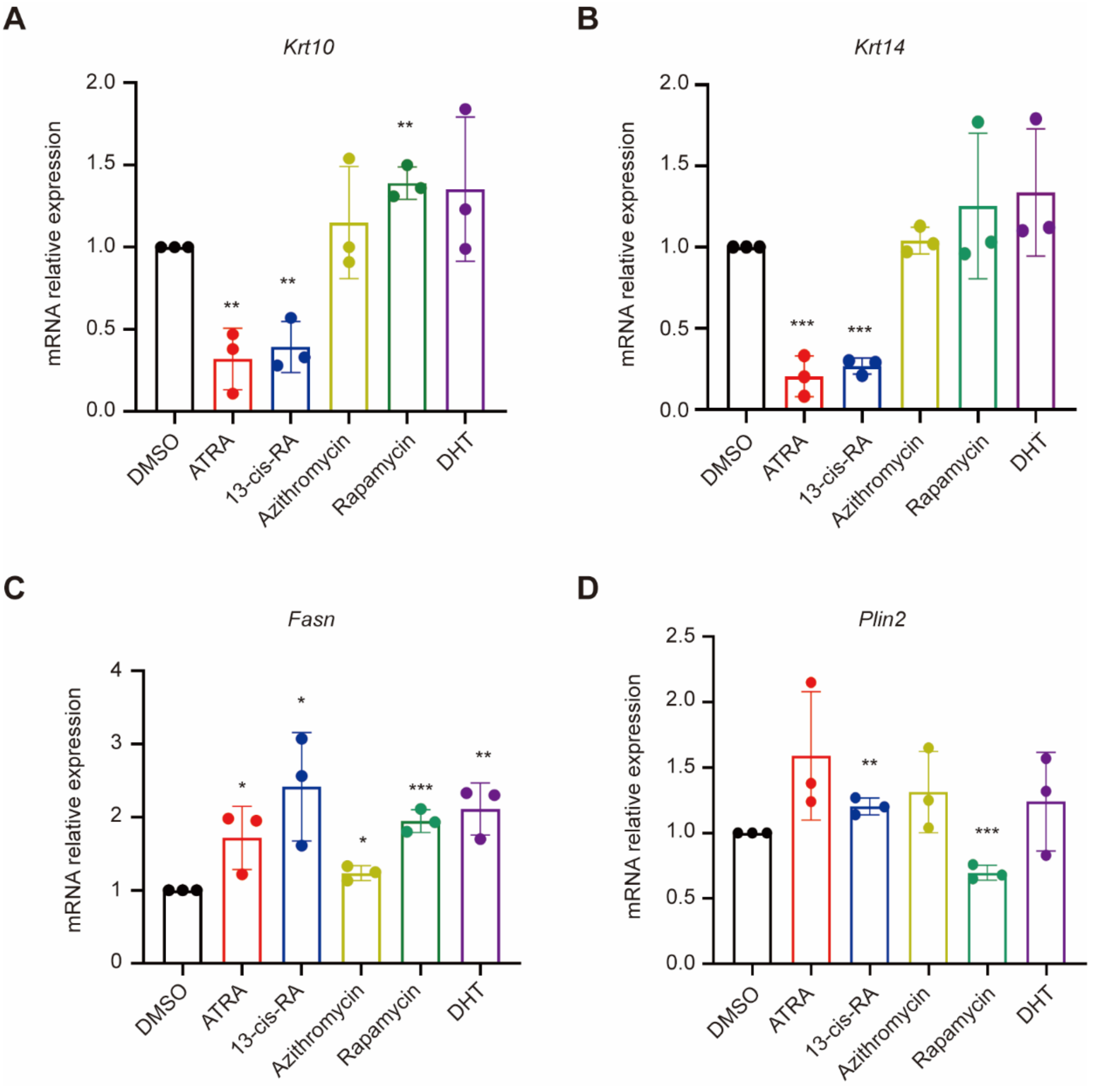


**Extended Data Fig. 4 | MG organoids can mimic the drug treatment responses of MGs *in vivo***

(A-D) Quantitative real time PCR (qRT-PCR) analysis of the expression level of *Krt5*(A), *Krt14*(B), *Fasn*(C) and *Plin2*(D) expression in MG organoids respectively incubated with ATRA, 13-cis-RA, Azithromycin, Rapamycin and DHT. Data were represented as mean ± SEM in 3 independent experiments. Unpaired two-tailed Student's *t* test: *$p$<0.05, ** $p$<0.01, ***$p$<0.001.

#### Extended Data Fig. 5

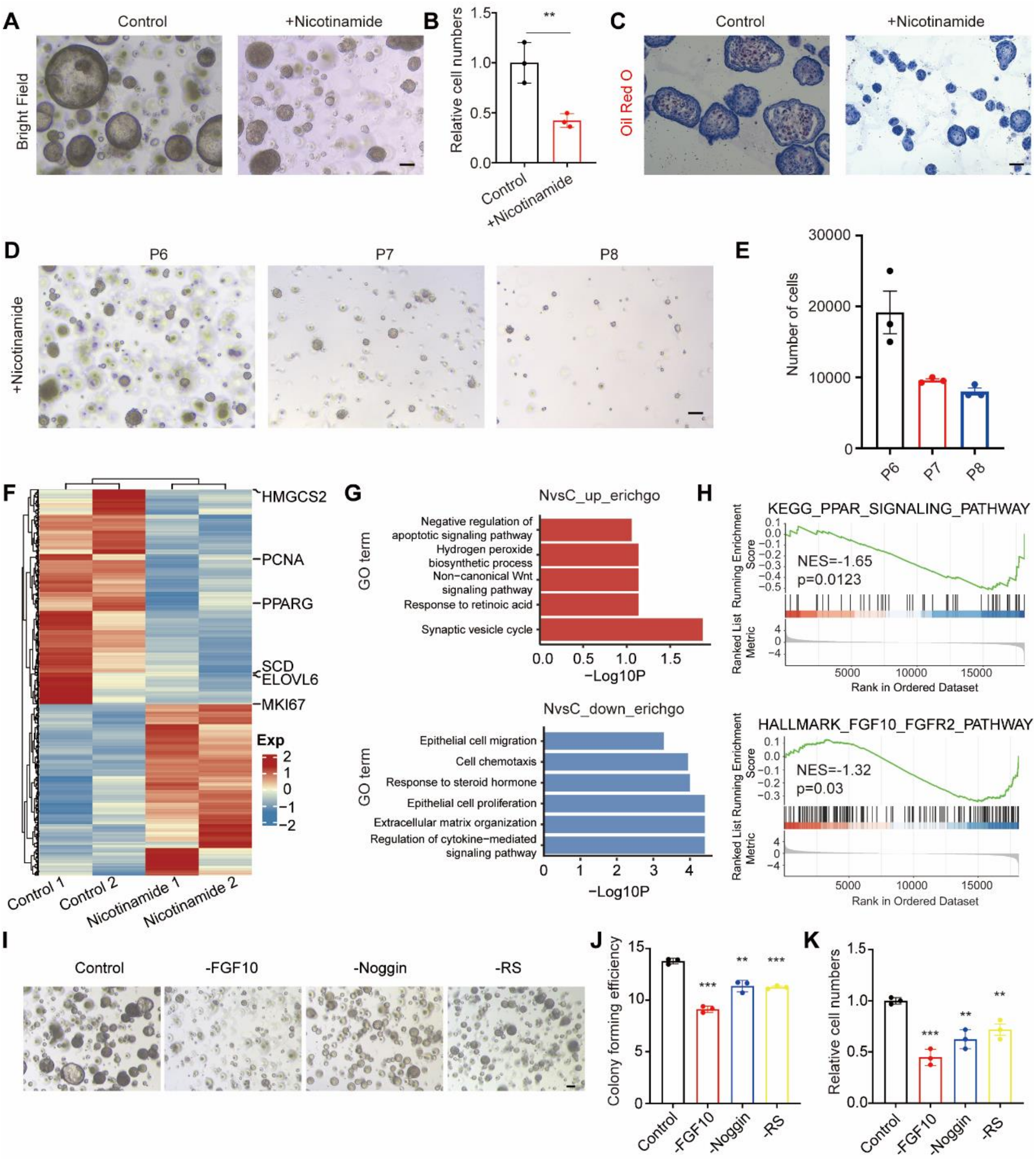


**Extended Data Fig. 5 | Nicotinamide inhibited the growth of hMGOs which required FGF10**

(A-B) Representative bright-field images(A) and relative cell numbers (B) of hMGOs in the culture medium with or without nicotinamide. Data were represented as mean ± SEM in 3 independent experiments. Unpaired two-tailed Student's *t* test: ** $p<0.01$. Scale bar, 100 μm.

(C) Oil Red O staining of lipid in hMGOs in culture medium with or without nicotinamide. Scale bar, 100 μm.

(D-E) Bright-field images (D)and cell numbers (E) of passage 6, 7 and 8 hMGOs cultured in medium with or without nicotinamide. Scale bar, 100 μm.

(F) Heatmap of the differentially expressed genes between the hMGOs under culture medium with or without nicotinamide.

(G) GO term enrichment analysis of the upregulated signaling pathway (above) and downregulated signaling pathway (below) in medium with nicotinamide compared to medium without nicotinamide.

(H) GSEA analysis of PPAR signaling pathway (above) and FGF10-FGFR2 signaling pathway (below) under the medium with nicotinamide added compared with that without nicotinamide added.

(I-K) The representative images (I), colony forming efficiency (J) and relative cell numbers (K) of the hMGOs outgrowth in 4 conditions (removing the FGF10, Noggin, Rspo-1 and control). Data were represented as mean ± SEM in 3 independent experiments. Unpaired two-tailed Student's *t* test: ** $p<0.01$, ***$p<0.001$. Scale bar, 100 μm.

#### Extended Data Fig. 6

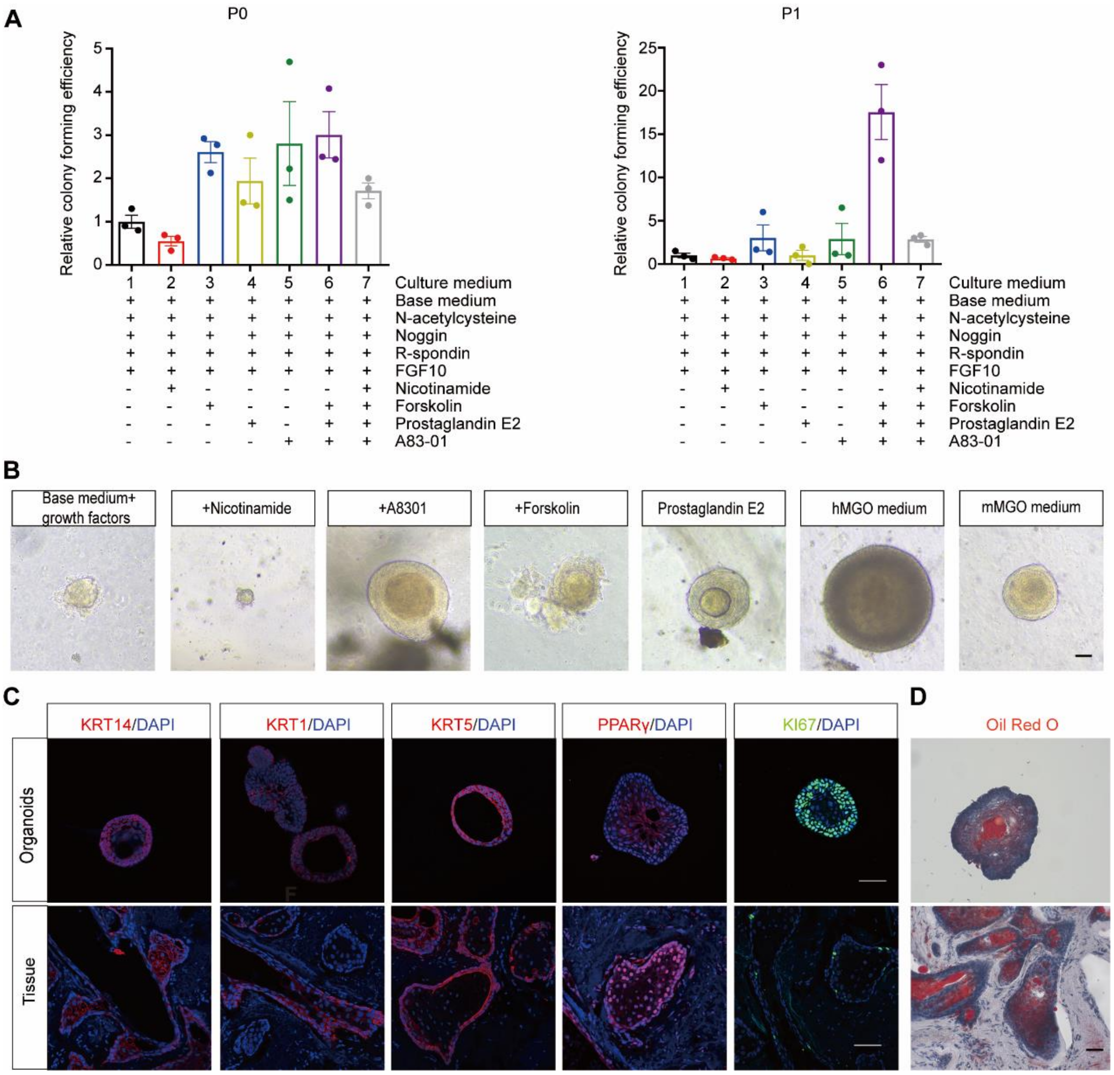


**Extended Data Fig. 6 | Identify the minimum composition of medium for hMGO culture**

(A) The relative colony forming efficiency of passage 0 and passage 1 organoids in different mediums. Scheme of 9 media compositions.

(B) Representative images of hMGOs outgrowth from primary tissue in different culture mediums. Scar bar: 50 μm.

(C) Immunofluorescent staining for the expression of MG epithelial cell marker (KRT14), ductal cell marker (KRT1), basal cell marker (KRT5), meibocyte marker (PPARγ) and the proliferation marker KI67 in hMGOs and tissue. Scar bar, 50 μm.

(D) Oil Red O staining of lipids in hMGOs and human MGs. Scale bar, 50 μm.

#### Extended Data Fig. 7

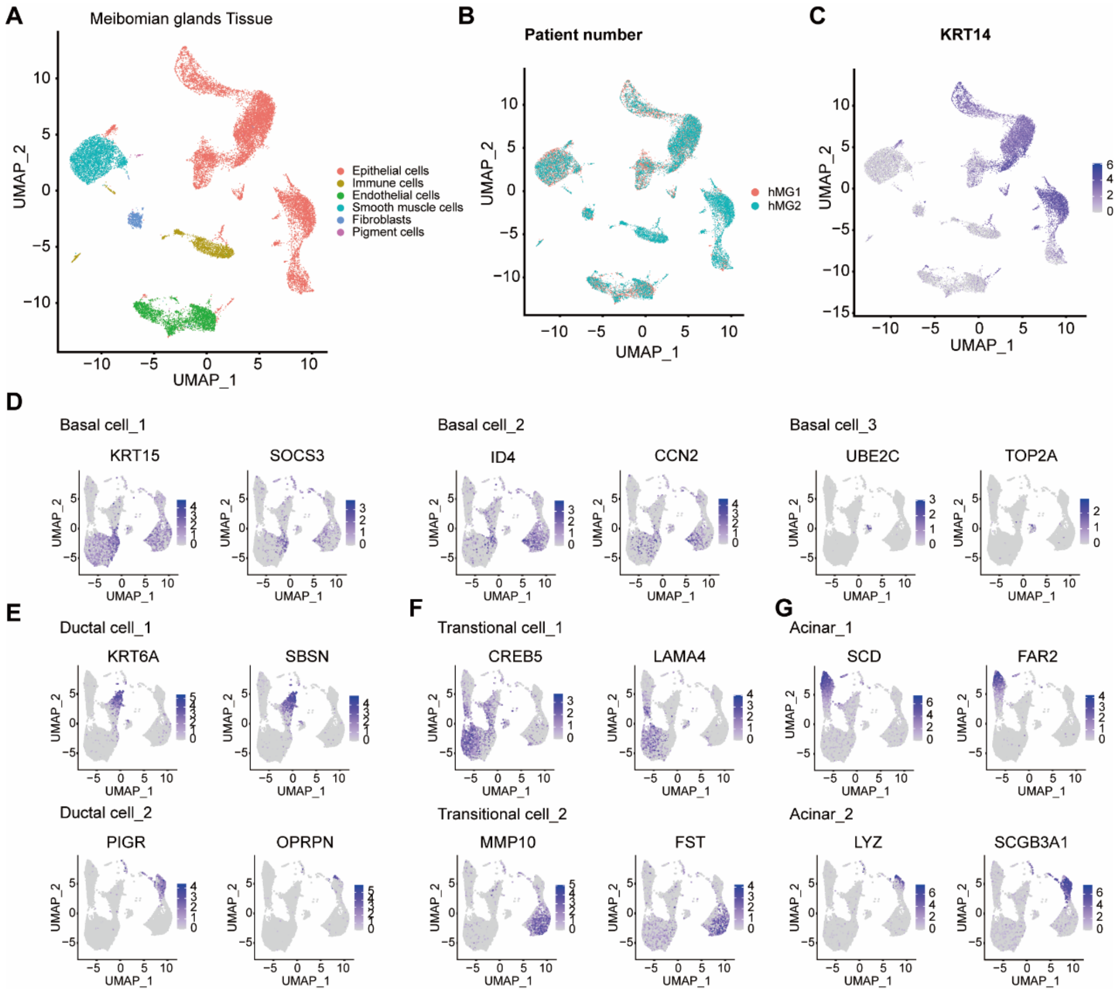


**Extended Data Fig. 7 | Dissection of cellular composition and heterogeneity by parallel single-cell atlas in human MGs and organoids**

(A) UMAP representation of six cell clusters identified in MG tissue (n=23,323).

(B) UMAP representation of MG cells from two individuals (tomato and cyan dots).

(C) The distribution of KRT14$^{+}$ cells in MGs tissue (n=13,684).

(D-G) The expression of cluster-specific genes in basal cells (D), ductal cells (E), transitional cells (F) and acinar cells(G), respectively.

#### Extended Data Fig. 8

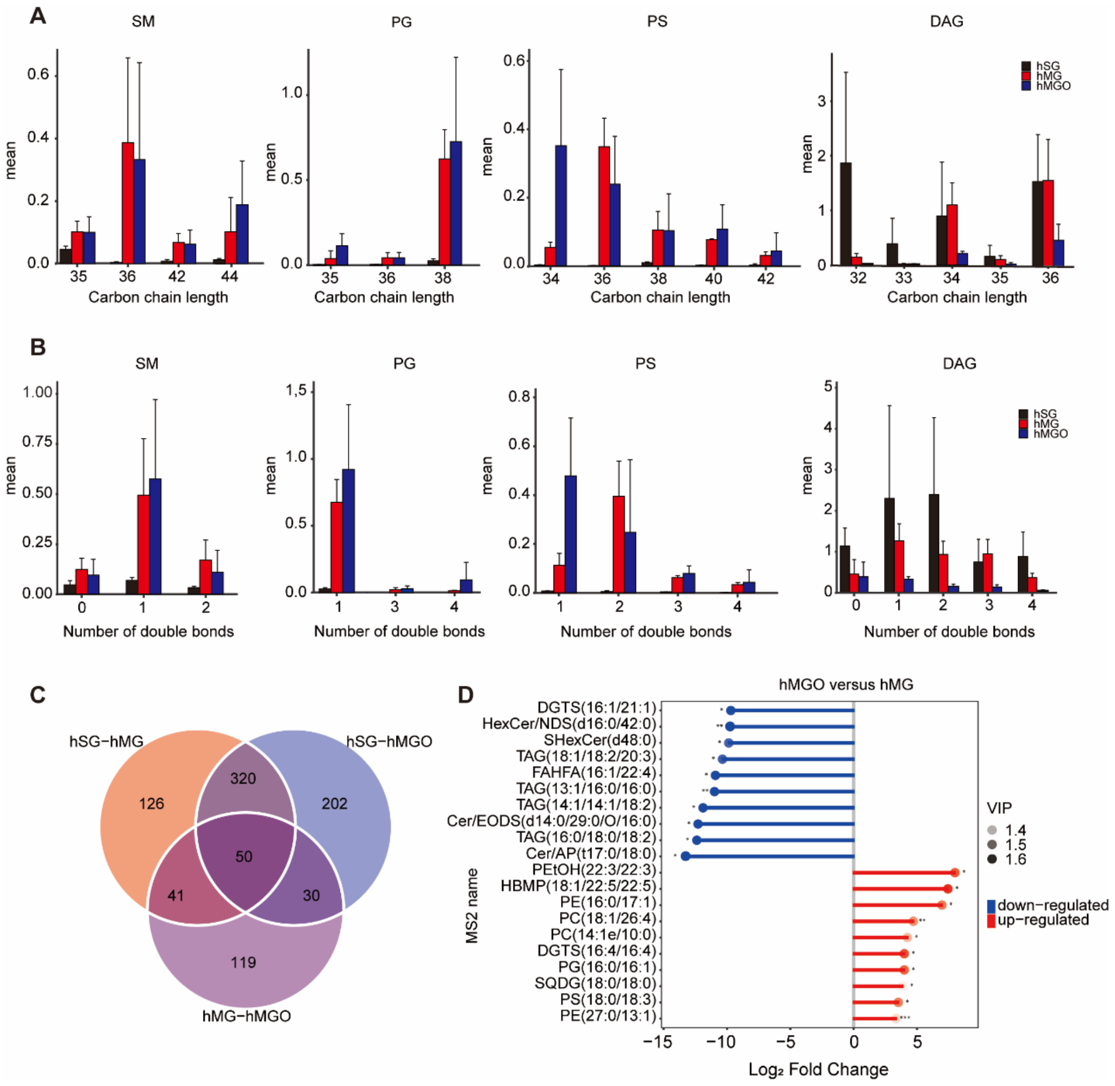


**Extended Data Fig. 8 | The non-targeted lipidomic analysis of hMGO and MG tissue**

(A) The carbon chain length analysis of sphingomyelins (SM), phosphatidylglycerols (PG), phosphatidylserines (PS) and diacylglycerols (DAG) in human MGs (hMG), human MG organoids (hMGO) and human sebaceous gland (hSG), respectively.

(B) The number of double bonds analysis of SM, PG, PS and DAG in hMG, hMGO and hSG.

(C) Differential lipids venn diagram of qualitative lipid species from nontargeted lipid profiling among 3 indicated groups. These lipids include 537 differential lipids between hSG and hMG, 602 differential lipids between hSG and hMGO, and 240 differential lipids between hMG and hMGO.

(D) Matchstick plot showing the top 10 up and down regulated lipids in hMGO compared with hMG.

### Extended Data Fig. 9

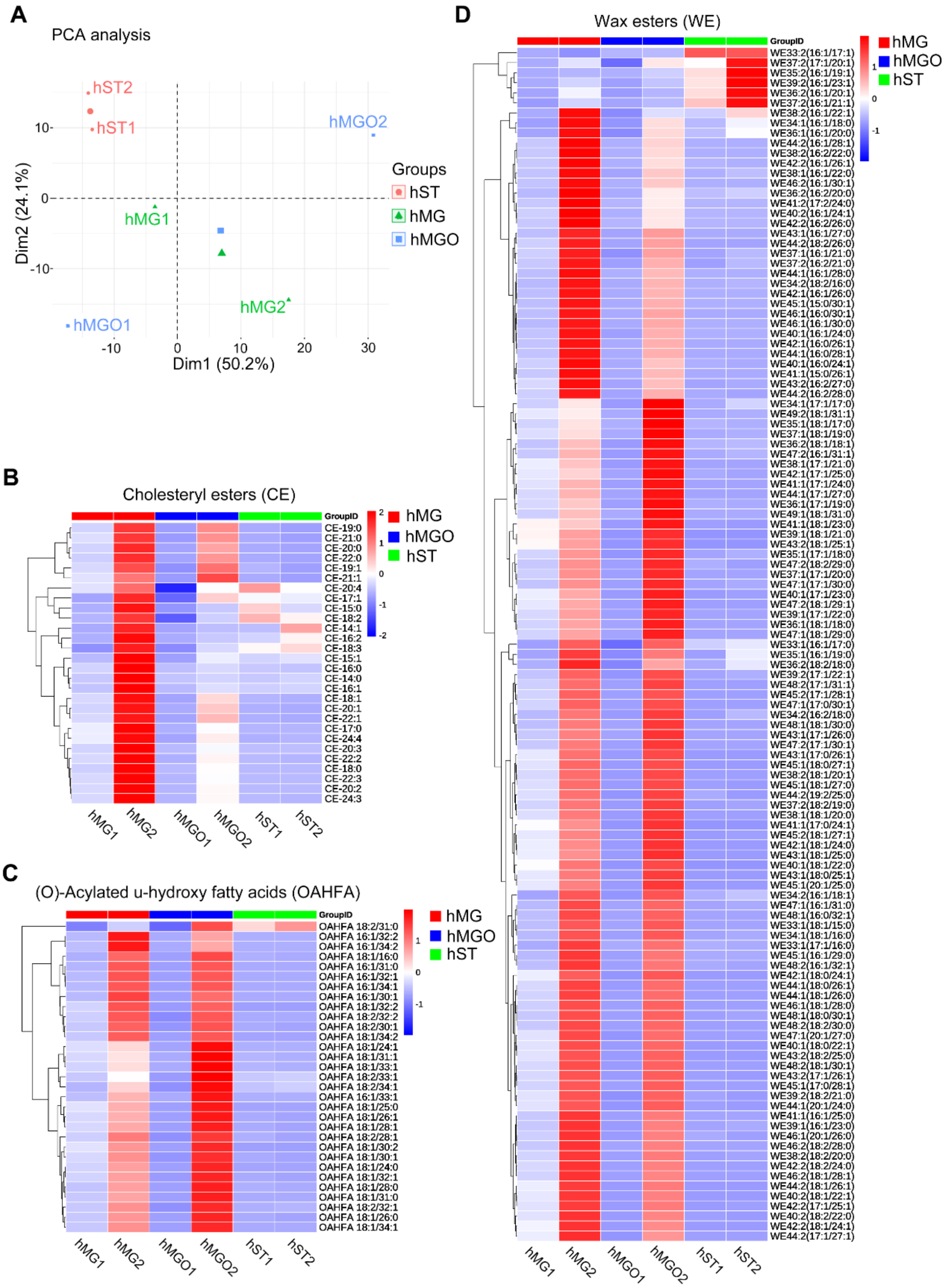


**Extended Data Fig. 9 | Targeting lipidomic analysis of human MG organoids and MG tissues**

(A) Principal component analysis (PCA) of lipids content between human MG tissue (hMG), human MG organoid (hMGO) and human periocular skin tissue (hST).

(B-D) Heatmap of cholesteryl esters (CE), (O)-Acylated u-hydroxy fatty acids (OAHFA) and wax esters (WE) between hMG, hMGO and hST.
